# Genetic control of gene expression and splicing in the developing human brain

**DOI:** 10.1101/471193

**Authors:** Rebecca L. Walker, Gokul Ramaswami, Christopher Hartl, Nicholas Mancuso, Michael J. Gandal, Luis de la Torre-Ubieta, Bogdan Pasaniuc, Jason L. Stein, Daniel H. Geschwind

## Abstract

Most genetic risk for human diseases lies within non-coding regions of the genome, which is predicted to regulate gene expression, often in a tissue and stage specific manner. This has motivated building of extensive eQTL resources to understand how human allelic variation affects gene expression and splicing throughout the body, focusing primarily on adult tissue. Given the importance of regulatory pathways during brain development, we characterize the genetic control of the developing human cerebral cortical transcriptome, including expression and splicing, in 201 mid-gestational human brains, to understand how common allelic variation affects gene regulation during development. We leverage expression and splice quantitative trait loci to identify genes and isoforms relevant to neuropsychiatric disorders and brain volume. These findings demonstrate genetic mechanisms by which early developmental events have a striking and widespread influence on adult anatomical and behavioral phenotypes, as well as the evolution of the human cerebral cortex.

**Highlights:**

- Genome wide map of human fetal brain eQTLs and sQTLs provides a new view of genetic control of expression and splicing.
- There is substantial contrast between genetic control of transcript regulation in mature versus developing brain.
- We identify novel regulatory regions specific to fetal brain development.
- Integration of eQTLs and GWAS reveals specific relationships between expression and disease risk for neuropsychiatric diseases and relevant human brain phenotypes.

## Introduction

Neurodevelopmental and neuropsychiatric disorders, such as autism spectrum disorder (ASD) and schizophrenia (SCZ), are highly heritable (Geschwind & Flint, 2015; Polderman et al., 2015) complex conditions, with hundreds of risk loci identified through large-scale genomic studies (Autism Spectrum Disorders Working Group of The Psychiatric Genomics, 2017; Gratten, Wray, Keller, & Visscher, 2014; Pardinas et al., 2018). However, the ability to interpret these susceptibility variants and their contributions to disease has been hampered because many of these variants fall in non-coding regions of the genome or in regions of high linkage disequilibrium, making it challenging to both identify causal mutations and their functional impact (Gandal, Leppa, Won, Parikshak, & Geschwind, 2016; Nica & Dermitzakis, 2013; Schaid, Chen, & Larson, 2018). Given the non-coding nature of the majority of these variants, as well as their enrichment in known regulatory regions (as inferred through chromatin accessibility (Cockerill, 2011; de la Torre-Ubieta et al., 2018), evolutionary conservation (Siepel et al., 2005), and signature histone modifications (Schaub, Boyle, Kundaje, Batzoglou, & Snyder, 2012; Visel, Rubin, & Pennacchio, 2009)), many of these variants likely function through the regulation of gene expression and splicing modulation (Y. I. Li et al., 2016; Maurano et al., 2012; Ward & Kellis, 2012). In this regard, multiple large-scale projects, including Roadmap Epigenomics, GTEx and Encode (Consortium, 2015; Ernst et al., 2011; Roadmap Epigenomics et al., 2015) have annotated regulatory regions across human tissues. However, little is known about how human allelic variation affects putative regulatory interactions during brain development, which is crucial for human higher cognition and brain evolution (Geschwind & Rakic, 2013; Nord, Pattabiraman, Visel, & Rubenstein, 2015; Ward & Kellis, 2012).

Expression quantitative trait loci (eQTL) analysis seeks to identify genetic loci that control changes in gene expression thereby annotating gene regulatory regions. Critically, eQTL relationships are highly dependent on cell type and developmental stage (Consortium, 2015; Flutre, Wen, Pritchard, & Stephens, 2013), which is consistent with transcriptional surveys of human brain development that show prominent temporal changes in expression (Colantuoni et al., 2011; H. J. Kang et al., 2011; Sunkin et al., 2013). Previous studies (Gilman et al., 2012; Gulsuner et al., 2013; Parikshak et al., 2013; Willsey et al., 2013) suggest that genetic disruption of these patterns, particularly during prenatal cortical development, leads to developmental and neuropsychiatric disease, highlighting the need to map regulatory variation within this critical time point. However, because developmental tissues are difficult to collect, all existing brain eQTL analyses, except for one study of small sample size (Jaffe et al., 2018), have been performed in adult brain (Consortium, 2015; Fromer et al., 2016; Ramasamy et al., 2014). Therefore, to dissect the functional genetic variation of neurodevelopmental and early onset neuropsychiatric diseases, characterized by phenotypes that likely originate in utero or early postnatal life (Hannon et al., 2016; Jaffe et al., 2016; Parikshak, Gandal, & Geschwind, 2015; Weinberger, 1987), we performed a well powered eQTL analysis in human cortex at mid-gestation, an epoch that captures crucial stages of neural progenitor proliferation and neurogenesis (M. B. Johnson et al., 2009; Silbereis, Pochareddy, Zhu, Li, & Sestan, 2016).

Here we present a map of eQTLs and spliceQTLs from fetal brain tissue to provide further understanding of mechanisms underlying transcriptional regulation and functional genetic variation in developing cerebral cortex. We have contrasted the genetic control of fetal and adult expression, which as expected do show substantial differences, as well as profiled the relationship between expression time points and disease risk. Overlap of eQTL and spliceQTL with GWAS identify new putative mechanisms missed by adult brain data sets, not only providing insights into genes and isoforms relevant to developmental risk factors, but also help to establish which aspects of disease risk are affecting neural progenitor proliferation and differentiation, as compared to later developmental processes like pruning, dendritic outgrowth, and myelination.

## Results

To identify genetic variants regulating gene expression in the developing brain, we performed high-throughput RNA sequencing and high-density genotyping at 2.5 million sites in a set of 233 fetal brains (**Figure 1**). After quality control and normalization of gene expression quantifications and genotype imputation into the 1000 Genomes Project phase 3 multi-ethnic reference panel (Methods, **Figure S1**; (Genomes Project et al., 2015), we obtained a starting dataset of 15,925 expressed genes (12,943 protein coding and 767 long noncoding RNAs) and 6.6 million autosomal single nucleotide polymorphisms (SNPs) from each individual. PCA-based (principle component analysis) analysis of ancestry (Methods) indicate that the donors in our study come from admixed ancestries of Mexican, European, African American, and Chinese descent (**Figure S2**). The resulting dataset is the first population level fetal brain expression dataset.

**Figure 1:**
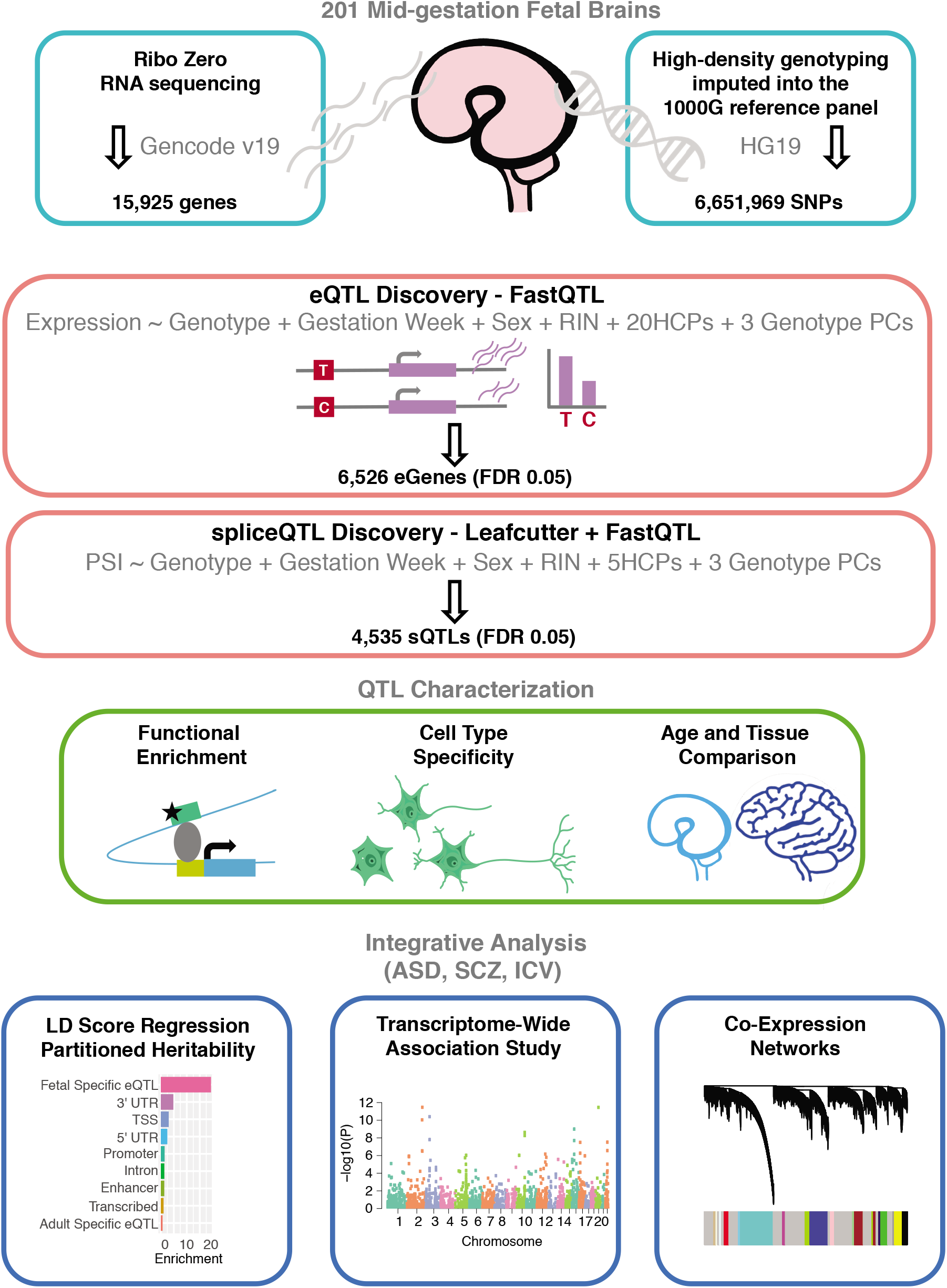
Study Design and Methodological Overview of QTL Analysis. RNA sequencing data from 233 fetal cortical samples was exhaustively QC’ed leaving data from 201 samples that was integrated with genome-wide imputation of common genetic variation based on high-density genotyping for expression and splicing QTL discovery. eQTL and sQTL were independently calculated using standard methods of covariate correction (Mostafavi et al., 2013) QTLs were characterized based on functional enrichment, cell type specificity, and compared to mature brain and non-brain tissue QTLs. We integrated these fetal brain QTLS with GWAS of neuropsychiatric disorders, performing LDSR and TWAS, as well as with regards to human brain phenotypes and gene co-expression networks to identify developmental disease risk and important biological processes under genetic control.

### Robust identification of fetal brain cis-eQTLs

We identified cis-eQTLs by testing all SNPs within a 1MB window from the transcription start site (TSS) of each gene using a permutation procedure implemented in FastQTL (Ongen, Buil, Brown, Dermitzakis, & Delaneau, 2016) while adjusting for known (RIN, sex, age, and genotype PCs) and inferred covariates (Methods, **Figure S3**), which have been shown to greatly increase sensitivity for cis-eQTL detection (H. M. Kang et al., 2008; Leek & Storey, 2007; Mostafavi et al., 2013). We identified 6,546 genes with a cis-eQTL at a 5% false discovery rate (FDR), hereafter referred to as eGenes, 82% of which corresponded to protein coding genes (Table S1). To compare with studies that did not perform a permutation procedure, we discover 893,813 nominal eQTLs at a 5% FDR. To identify additional, independent cis-eQTLs for each eGene, we reran the analysis while conditioning on the primary eSNP, identifying an additional 1,416 secondary eQTLs for a total number of 7,962 eQTL identified, which represents nearly 45% of expressed genes (Methods). To ensure that the eQTLs identified are not driven by ancestry differences within our dataset, we ran eQTL analysis using two alternative methods: EMMAX, which controls for population stratification using a genetic relationship matrix (H. M. Kang et al., 2008) and METAL, which performs meta-analysis of the top SNP per gene across subpopulations of the samples (Willer, Li, & Abecasis, 2010). Both methods demonstrated high reproducibility of eQTLs identified by fastQTL, which indicates that ancestry differences are not driving the eQTL signal (**Figure S5**; Methods).

### eQTLs mark functionally active regulatory regions

Positional enrichment analyses of the most significant SNP (eSNP) at each eQTL (primary and secondary) shows that over 20% of significant eQTLs cluster with 10kb of the TSS of its target gene (**Figure 2B and 2C**), concordant with previous studies showing that promoter variants have a large influence on cognate gene expression (Y. Kim et al., 2014; Stranger et al., 2007; Strunz et al., 2018). The overall distribution of eQTL signal is consistent with previous work (Consortium, 2015; Veyrieras et al., 2008), with 70% of significant eSNPs located within +−100kb of the TSS, as well as a slight upstream bias (~56%) of significant eQTLs (**Figure 2B**). We reasoned that eQTL should be highly overlapping with putative regulatory regions defined by other methods and sought to test this as an external validation (**Figure 2A**). Indeed, we find eQTLs are significantly enriched within regions of open chromatin identified by ATAC-seq from developing brain (OR=4.42, p-value < 2.2e-16) (**Figure 2D**; (de la Torre-Ubieta et al., 2018). Distal eQTLs (> 10Kb from TSS) are also enriched within 3D chromatin conformation contacts at the same stage of brain development (OR=2.96, p-value < 2.2e-16) (**Figure 2E**; (Won et al., 2016) strengthening confidence that eSNPs are in accessible regions where transcription factors preferentially bind and distal eSNPs regulate their associated target eGenes (Methods).

**Figure 2:**
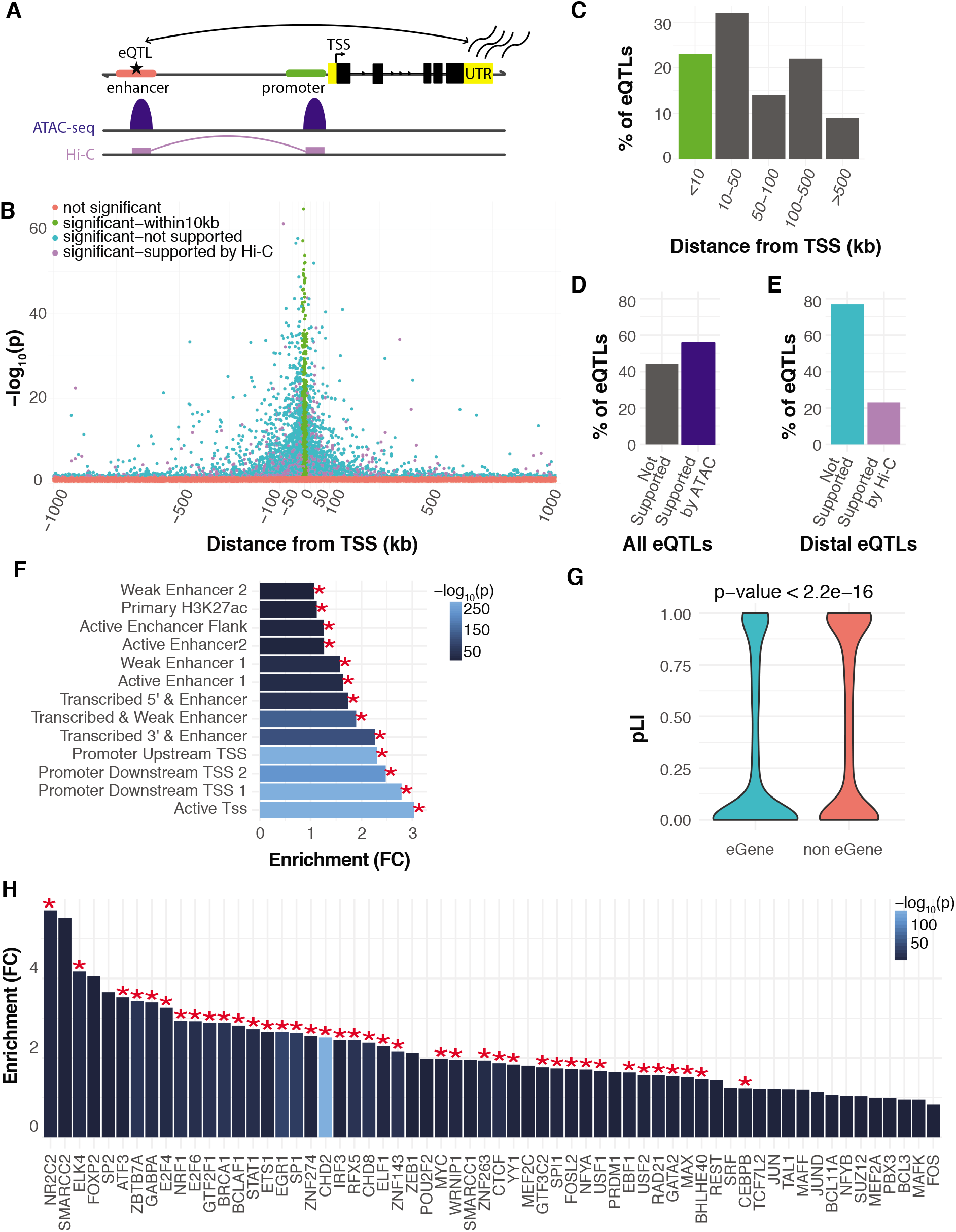
Characterization of cis-eQTLs. **(A)** A schematic showing chromatin accessibility (ATAC-seq) and chromatin interactions (Hi-C) are used to support eQTLs in linking genes to distal regulatory elements. **(B)** Position of primary and secondary eQTLs, annotated by their color and position as depicted in the key. eSNPs are shown in relation to the eGene transcription start site (TSS) as defined in Genecode v19. Significant eQTLs are annotated based on support from fetal brain chromatin contacts defined by Hi-C (Methods) and distance from the TSS. **(C)** Distribution of all primary and secondary cis-eQTLs; most lie > 10kb 3’ or 5’ to the TSS. eQTLs that lie within 10kb of the TSS are colored green, as depicted in (A). **(D)** Fraction of eQTLs with fetal brain ATAC support. **(E)** Fraction of distal eQTLs (eSNP is further than 10kb away from the TSS) with fetal brain Hi-C support. eQTLs that show Hi-C support are colored purple, while those not supported by Hi-C are colored blue, as depicted in (A). **(F)** eGenes are more tolerant to loss of function, as compared with those genes expressed in fetal brain without significant cis-eQTL. **(G)** Fold enrichment of eSNPs by their distribution within specific fetal brain chromatin states, which are listed on the Y axis (ChromHMM annotations; Methods). eQTL are in active states, most prominently in promotors and nearby cis-regions, as expected. **(H)** Fold enrichments of eSNPs in experimentally discovered transcription factor binding sites from (Arbiza et al., 2013) characterize the intersection of transcriptional control elements with human genetic variation.

To further characterize the identified eQTLs, for each eGene, we annotated the most significant variant with chromatin state predictions from fetal brain tissue from the Roadmap Epigenetics Consortium (Roadmap Epigenomics et al., 2015), using GREGOR (Schmidt et al., 2015) Methods) to test for enrichment of eQTLs among the 25 chromatin states. We observed that eSNPs were most significantly enriched in transcription start sites, promoters, and transcribed regulatory promoter/enhancers (**Figure 2F**, full model **Figure S6A**). Enrichment in these regulatory regions provides further evidence, in addition to the ATAC and Hi-C, that discovered eQTLs fall in defined regions of the genome functionally relevant to gene expression and its regulation, as would be predicted (Ward & Kellis, 2012).

### Functional characterization of eQTLs reveals mutation tolerance and regulatory drivers

Since changes in constrained genes expressed in the developing brain are likely to have a higher impact on fecundity than those that are not highly constrained (Samocha et al., 2014), we reasoned that genes perturbed by regulatory variation would also be more tolerant to protein-disrupting variation (Lek et al., 2016). Indeed, comparing genes harboring an eQTL to genes that do not have a significant eQTL, we find that eGenes are more tolerable to loss of function mutations and less constrained (Wilcoxon rank sum test p-value<2.2×10 −16; **Figure 2G**).

To identify putative transcription factors regulating the expression of genes within the fetal brain, we examined whether eSNPs identified are enriched in transcription factor and DNA binding protein (DBP) binding sites, using multiple cell types from the ENCODE project (Methods; (Arbiza et al., 2013). We found significant enrichments for 39 transcription factors and DBPs binding sites (**Figure 2H**), many with prominent known roles in brain development, including FoxP2, POU2F2, ELK4, NRF1, CHD8 (Bestman, Huang, Lee-Osbourne, Cheung, & Cline, 2015; Durak et al., 2016; Eising et al., 2018; Lai, Fisher, Hurst, Vargha-Khadem, & Monaco, 2001; Ojeda et al., 1999; Preciados, Yoo, & Roy, 2016). Many of these genes have previously been associated with developmental disorders. The most significantly enriched protein was CHD2 ( FC= 2.5, p-value= 2.5×10^-143^), a chromatin remodeling protein that shows the highest expression during early developmental periods in fetal cerebral cortex (Sunkin et al., 2013).

Similar to several of the other transcription factors including CHD8, FoxP2 and SMARCC2, haploinsufficiency in CHD2 has dramatic pathogenic consequences, and has been associated with risk for autism spectrum disorder (ASD), developmental delay, intellectual disability, seizures and epilepsy, similar to CHD8 (Carvill et al., 2013; Suls et al., 2013).

### Robust identification of fetal brain splice QTLs

Effects of genetic variation on RNA splicing are known to contribute to complex disease risk, comparably, if not more than eQTL (Y. I. Li et al., 2016). Since there has been no systematic analysis of splicing in human fetal brain development, we reasoned that such analysis would be critical to further the understanding of the link between genetic variation and disease. We quantified intron clusters using Leafcutter (Y. I. Li et al., 2018), identifying 92,449 intronic excision clusters which mapped to 10,926 genes. A major strength of Leafcutter is that it is annotation free, allowing for alternative exon discovery, which is especially important in the context of brain isoforms, for which annotations are known to be incomplete. Per sample intron abundances (PSI, percent spliced in) were then used as a quantitative molecular trait for local spliceQTL (sQTL) discovery in fastQTL (Ongen et al., 2016); Methods, **Figure S3**), identifying a total of 4,635 significant sQTLs (FDR < .05) in 2,132 genes (sGenes; **Table S2, Figure 3A**).

**Figure 3:**
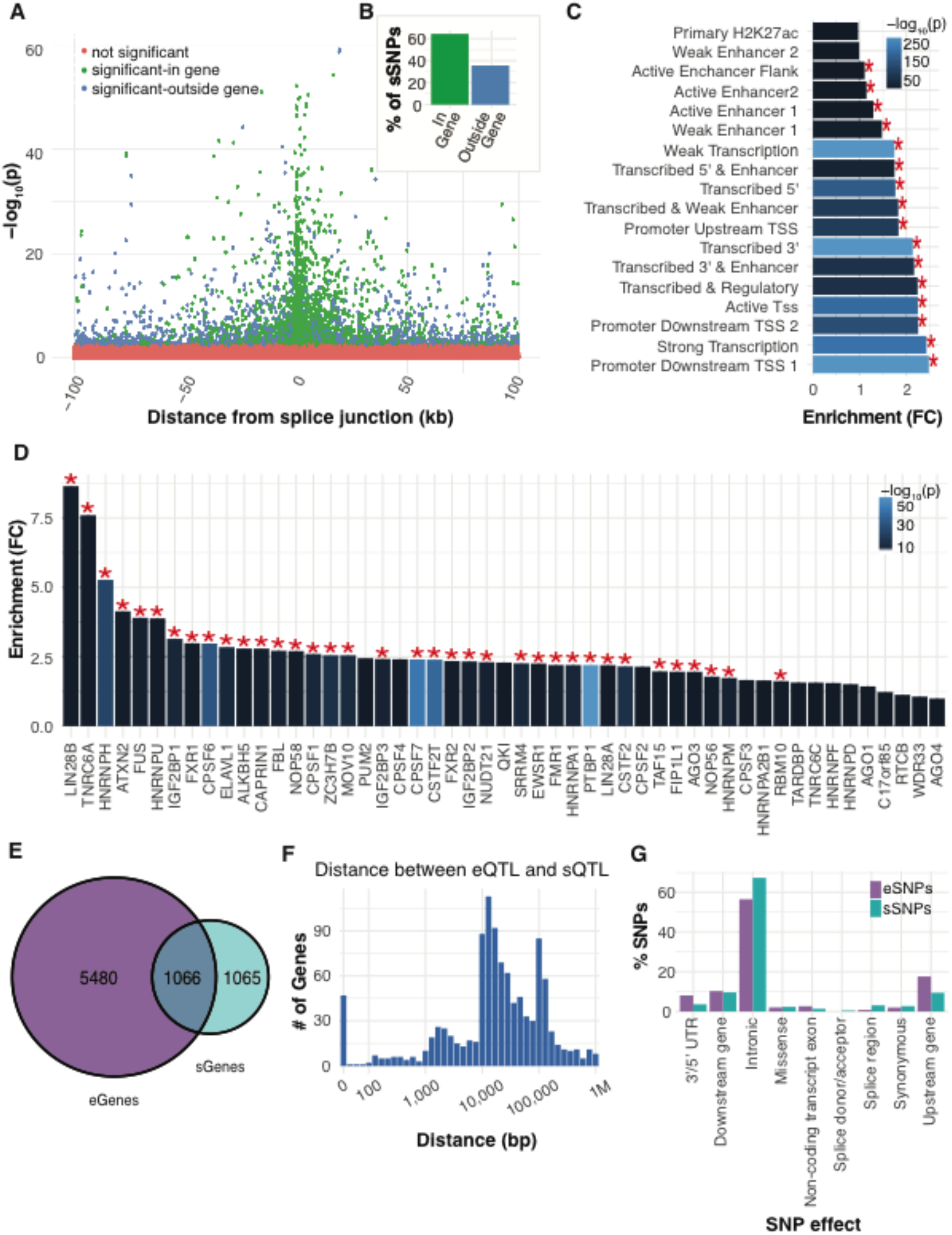
Characterization of sQTLs and Comparison of sQTLs to eQTLs. **(A)** Position of sQTLs in relation to the splice junction. Significant sQTLs are annotated based on whether the sSNP lies within (green) or outside (blue) of the corresponding sGene. **(B)** Fraction of sQTLs where the sSNP lies within vs outside its sGene. **(C)** Fold enrichment of sSNPs by their distribution within specific fetal brain chromatin states, which are depicted on the Y axis (ChromHMM annotations; Methods). sSNPs are most prominently found within promoters and transcribed regions. **(D)** Fold enrichments of sSNPs in experimentally discovered RNA binding protein binding sites from the CLIPdb database (Y. C. Yang et al., 2015) (Methods). **(E)** Venn diagram showing the overlap of genes containing an eQTL (6,546) vs sQTL (2,132). **(F)** Distribution of the distance in base pairs between eSNP and sSNP for genes harboring both. **(G)** Relative proportions of predicted SNP effects (VEP) between eSNPs and sSNPs.

### Genomic features of sQTLs distinguish them from eQTL

Positional enrichment of the top SNP per spliceQTL (sSNP) shows clustering around the splice junction, with 42% of sSNPs within 10 kb of the splice junction (**Figure 3A**), demonstrating that variants proximal to splicing junctions have a large effect. In contrast to eQTL, the majority of sSNPs (64%) lie within the gene body (**Figure 3B**), consistent with expectations (Y. I. Li et al., 2016). Splice QTLs were most strongly enriched in promoters and transcribed regions of the Roadmap Epigenetics Consortium chromatin states from fetal brain tissue (**Figure 3C**, full model **Figure S6B**), consistent with regions functionally relevant to gene splicing mechanisms (Lappalainen et al., 2013; Takata, Matsumoto, & Kato, 2017).

### Identifying drivers of fetal brain RNA-splicing

To identify putative splicing regulators in the human fetal brain, we checked for sSNP enrichment in RNA binding protein (RBP) binding sites using the CLIPdb database of publicly available cross-linking immunoprecipitation (CLIP)-seq datasets from 51 RBPs (Y. C. Yang et al., 2015). We found binding sites for 36 of the RBPs to be significantly enriched among sSNPs (**Figure 3D**; Methods), most of which do not have well characterized roles in CNS function, especially during neural development. Among the significant RBPs with the sSNP enrichment with known roles in neurodevelopment are HNRNPH, ATXN2, and SRRM4. HNRNPH is involved in many aspects of neurodevelopment from alternative splicing of TRF2, which is implicated in neuronal differentiation (Grammatikakis et al., 2016), to oligodendrocyte differentiation (Wang, Dimova, & Cambi, 2007). ATXN2 plays roles in proliferation and cell growth with known mutations in the gene itself causing neurodevelopmental phenotypes, from spinocerebellar ataxia, ALS and schizophrenia (Almaguer-Mederos et al., 2018; Y. E. Kim et al., 2018; F. Zhang et al., 2014). SRRM4 is a splicing factor known to regulate alternative exons with increased neural inclusion and including many microexons, with loss of function in mice linked to numerous neurodevelopmental deficiencies (Quesnel-Vallieres, Irimia, Cordes, & Blencowe, 2015).

### eQTL and sQTL are distinct, but overlapping

To assess sharing of eQTL and sQTLs, we next analyzed the overlap of genes harboring a significant eQTL compared to those with sQTLs. We found that about half of all genes with a spliceQTL were also an eGene (1,066 genes out of the 6,546 eGenes; **Figure 3E**). To determine the extent in which eQTL and sQTL might have overlapping effects, we performed a threshold free comparison of significant sQTLs from Leafcutter (SNP and splice junction mapped to genes) with eQTL summary statistics to estimate the proportion of non-null associations among the eQTLs (Storey & Tibshirani, 2003); Methods). We find that 68% of sQTLs also affect total levels of gene expression; however, only 19% of eQTLs affect splice-junction usage, suggesting mostly independent regulation of expression and splicing events, as has been observed in other tissues (Y. I. Li et al., 2016). This independence of expression and splicing regulation is also observed when comparing eSNPs and sSNPs for the same gene, as most of the tagged regulatory regions for expression and splicing of the same gene are distinct, as evidenced by the large distances between eSNP and sSNP (**Fig 3**). At the same time, we note that there are a minority of SNPs that affect both total expression and splicing, as evidenced by the 47 genes with directly overlapping sQTL and eQTL (**Figure 3F; Table S3**). We also used Ensembl’s Variant Effect Predictor (VEP) (McLaren et al., 2016); Methods) which annotates SNP function, to compare the distributions of our splicing and expression related variants, finding very similar results. For example, we observe more sSNPs identified as an intronic variant than eSNPs, and more eSNPs being identified as upstream gene variants (**Figure 3G**).

### Tissue specificity corresponds to eQTL effect size

To examine eQTL sharing among tissue types, we next examined the correlation of effect sizes of fetal brain eQTLs (**Figure S7**; (Nica et al., 2011) to those from 48 different tissue types from the Genotype-Tissue Expression Consortium (GTEx) (Methods; (Consortium et al., 2017). Fetal eQTLs found in one or more GTEx tissue types showed significantly lower effect sizes (p-value <2.2×10 −16) than those that were fetal specific, suggesting eQTLs active globally across tissue types are more constrained, whereas fetal brain specific eQTLs have greater magnitude of effect sizes (**Figure 4A**). That fetal brain specific eQTLs are less constrained than broad eQTL is also supported by comparison of a measure of tolerance to loss of function mutations, pLI (Lek et al., 2016), between eGenes specific to fetal brain or shared between one or more GTEx tissue types (**Figure 4B**). To assess fetal eQTL sharing between each tissue, we correlated the effect sizes of all significant fetal eQTLs to the same eQTL SNP-gene pair across all GTEx tissues (Methods). We observe the strongest Spearman’s *ρ* correlations of fetal brain eQTLs with adult brain tissue and proliferative epithelial containing tissues (**Figure 4C**)

**Figure 4:**
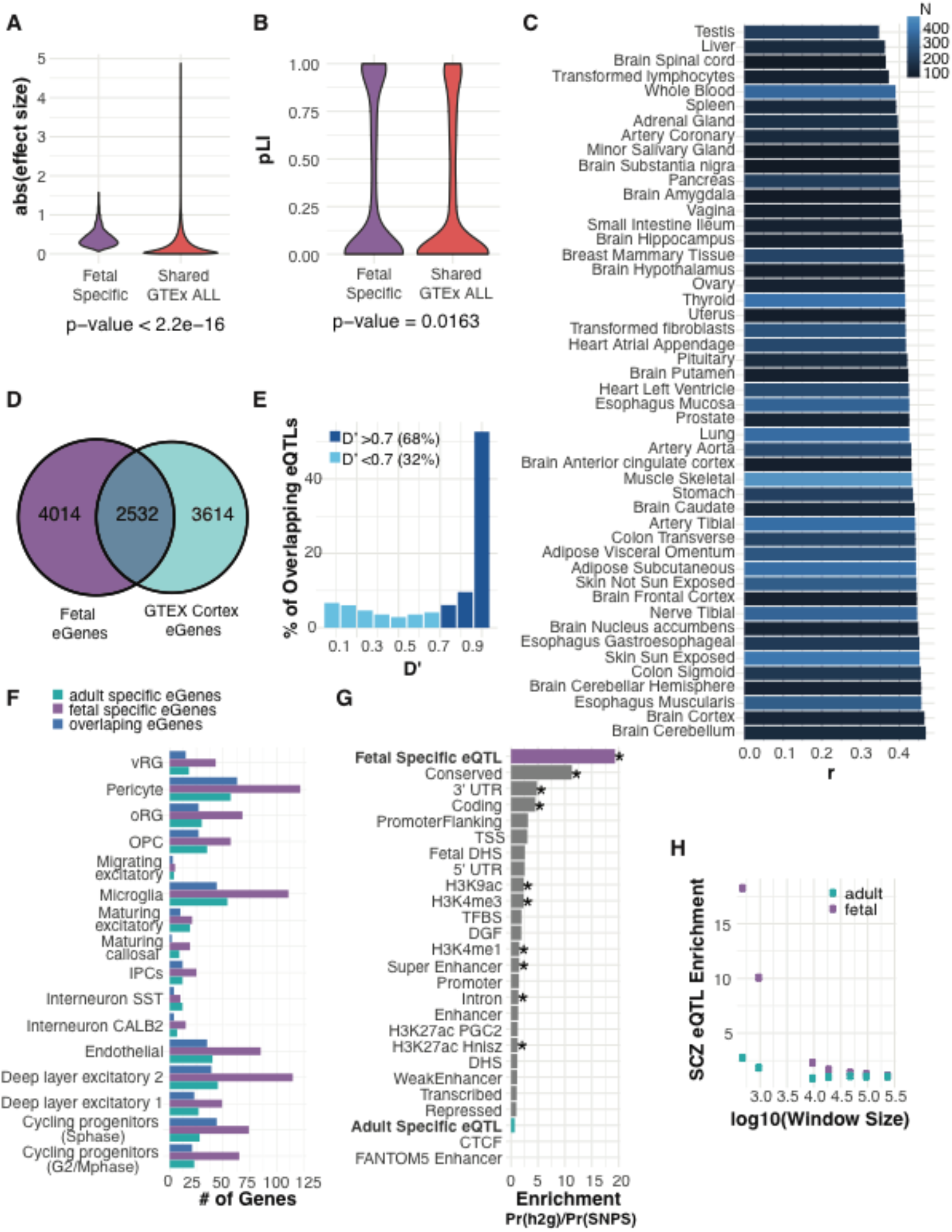
Age specificity of Brain eQTLs. **(A)** Distribution of effect size for all fetal eQTLs, fetal-specific eQTLs vs eQTLs identified in any GTEx tissue. **(B)** Distribution of pLI scores of loss-of-function intolerance for all fetal eQTLs, including fetal-specific eQTLs vs eQTLs identified in any GTEx tissue. **(C)** Effect size correlations (Spearman’s *ρ*) between significant nominal fetal eQTLs (absolute value of beta) and corresponding eQTL among GTEx v7 tissues. Bars are colored by the GTEx eQTL dataset sample size. The highest correlations are seen with brain and proliferative, epithelial tissues. **(D)** Venn Diagram comparing genes with an eQTL discovered in fetal cortical tissue vs adult cortical tissue from GTEx (Consortium et al., 2017); Methods). **(E)** Distribution of linkage disequilibrium (D’) between eSNPs of genes that overlap between fetal and adult datasets. This comparison shows that the majority of eQTLs are tagging the same region, even when the eSNP differs between the adult eQTL and fetal eQTL. **(F)** The number of fetal single cell cluster-marker genes (Methods; significantly differentially expressed between all other clusters with a log(FC)>0.2) containing an eQTL in fetal-specific eQTL data, adult-specific eQTL data, and those overlapping both datasets. Fetal = fetal specific; adult = adult specific; both = overlapping and found in both fetal and adult. **(G)** LD score regression annotation enrichments where annotations of fetal eQTLs and adult eQTLs were added to the baseline annotations by creating a 500bp window around the eSNP and removing any region observed in both fetal and adult annotations. An asterisk indicates significance at P<0.05 after Bonferroni correction for all annotation categories tested. It is notable that the distribution of shared QTL essentially mirrors the adult, while the fetal-specific show a distinct distribution, especially within primarily fetal cell types. **(H)** LD score regression enrichment of the fetal specific annotations and adult specific annotations with varying window size around the eSNP. Enrichment of SCZ in fetal eQTL annotations is consistent across window sizes.

### Fetal eQTL are distinct from those in Adult

Another central question is related to developmental stage: how do fetal brain eQTLs compare with adult brain eQTLs? To address this, we first compared genes that harbor an eQTL in our dataset to that of GTEx adult cortex eQTLs (N=136 individuals, eGenes=6,146), a dataset of similar sample size, experimental design, and number of significant eQTLs (Consortium et al., 2017). We found 2,532 eGenes that overlapped between fetal and adult, accounting for a little over one third of both datasets (**Figure 4D**). Of the eGenes that overlapped, we examined how many eSNPs were tagging the same region by calculating the LD between the top primary SNPs in the fetal and adult data in the 1000 Genomes Project phase 3 multi-ethnic reference panel (Methods; Genomes Project et al., 2015). We found 68% of overlapping eQTLs tag the same regulatory region, even if the top SNP differed (**Figure 4E**). To further examine the differences in eGenes at the different developmental time points, eGenes were annotated based on fetal cell type markers, identified through differential expression of single cell sequencing identified cell clusters (Methods; Polioudakis et al., 2018). We find many more fetal eGenes are in fact fetal cell type markers compared to eGenes that either overlap with adult, or are adult specific (**Figure 4F**), which is expected given the expected cellular composition of each epoch.

### Integrating GWAS signal in the context of development

A major utility of eQTL studies is to provide functional annotation of disease-associated variants, largely identified from GWAS. Recent studies have shown heritability for complex disorders are disproportionately enriched in functional categories such as conserved regions and enhancers (Finucane et al., 2015). Stratified LD score regression can be layered on top of functional annotations to estimate the regulatory regions’ contributions to common genetic disease risk as assessed by GWAS. Previous work based on active chromatin during fetal development had suggested that liability for SCZ was significantly enriched in fetal brain, especially in the progenitor containing zones (de la Torre-Ubieta et al., 2018). To test this, we created eQTL annotation regions for fetal cortex and adult cortex by considering a 500bp window (+-250) around each eSNP and removed overlapping annotation windows, resulting in a comparable number of annotations for each epoch (6,163 fetal annotations, 5,690 adult annotations, Methods). Partitioned GWAS heritability for SCZ shows fetal brain eQTLs to be the highest enriched among all significant functional categories, with 0.2% of SNPs explaining an estimated 4% of SNP heritability (P = 9.1×10^-4^ for enrichment), and adult brain eQTLs not reaching significance (**Figure 4G**), suggesting that fetal brain regulatory regions harbor a greater proportion of SCZ risk variant than adult. To test the robustness of this finding, we explored different window sizes around each eGene, removing any window that overlapped between fetal and adult annotations, and saw a consistent significant enrichment for fetal annotations over adult for SCZ GWAS loci (**Figure 4H**).

### eQTL within the context of transcriptional networks

We reasoned that our sample size, which is several-fold larger than any previous gene expression study of fetal brain (H. J. Kang et al., 2011; Parikshak et al., 2013), would permit identification of robust sets of co-expressed genes. We applied robust weighted gene coexpression network analysis (B. Zhang & Horvath, 2005) to construct transcriptional networks, identifying 19 modules (labeled by color) of co-expressed genes during mid-gestation cortical development (**Figure 5A, Table S4**). The modules identified represent genes that correspond to distinct biological functions defined through shared gene ontology (GO) enrichments and cell type markers within the module (**Figure 5B and 5C**, Methods). Six of these modules are enriched for specific brain cell types or brain-relevant ontological terms: turquoise (fetal mitotic progenitors, cell division), red (fetal mitotic progenitors, outer radial glia, splicing), yellow (fetal neurons, splicing), blue (fetal neuron, axon guidance), greenyellow (adult neuron, synaptic transmission, neuron projection development), and brown (adult neuron, CA2+ transport). All of these modules show high preservation in an independent RNA-seq dataset of cortical development from 8 post conception weeks to 12 months after birth (Parikshak et al., 2013; Sunkin et al., 2013), supporting the notion that this window in mid-gestation, by containing a full range of the major cell types, from proliferating progenitors and post mitotic migrating to post migratory neurons, captures a substantial portion of the biological processes occurring during brain development (Polioudakis et al., 2018; Pollen et al., 2015).

**Figure 5:**
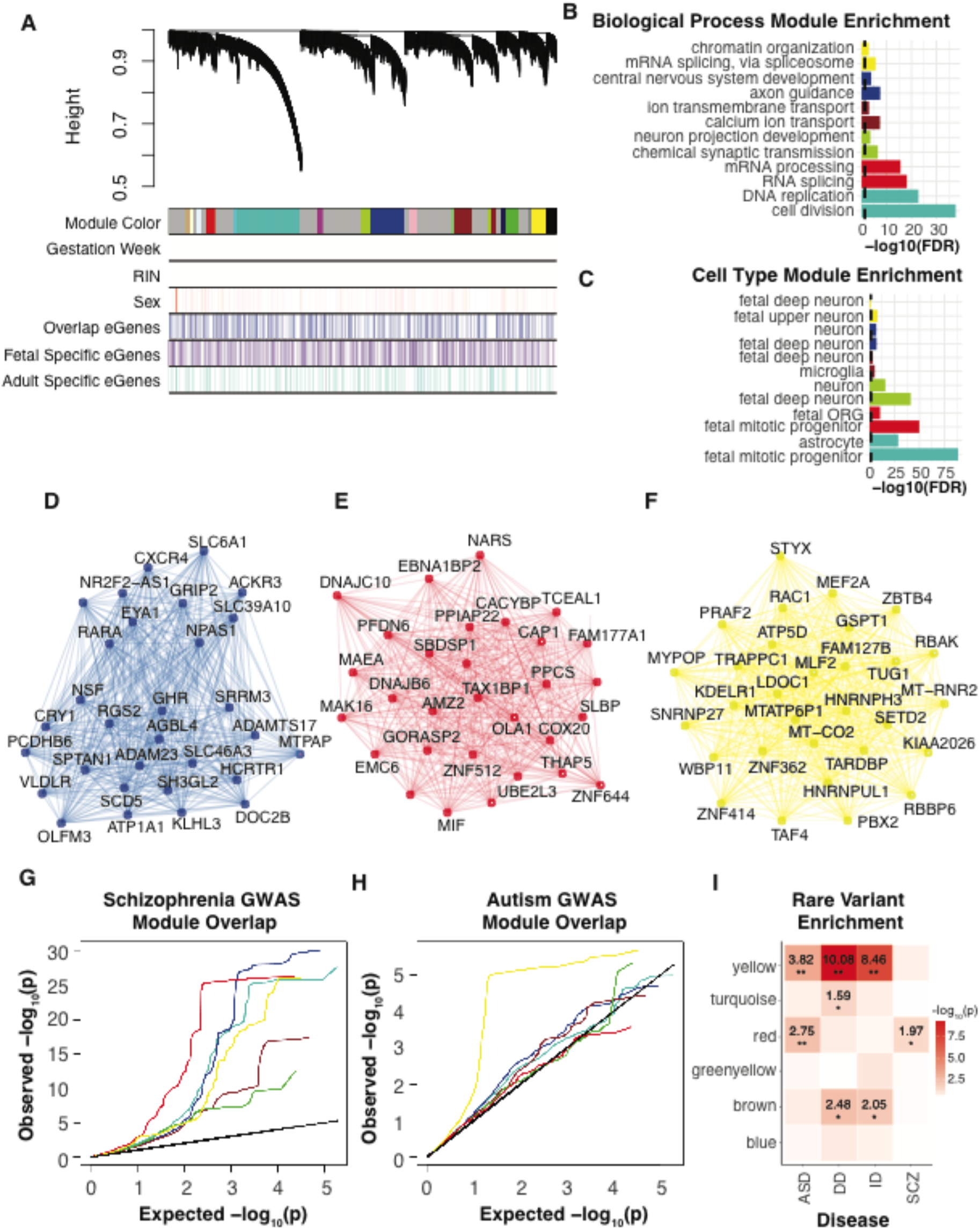
Fetal Brain Co-expression Networks. **(A)** Network analysis dendrogram based on hierarchical clustering of all genes by their topological overlap, identifies 19 developmental modules (Methods). Colored bars below the dendrogram show module membership, expression covariates (Gestation week, RIN, and Sex), and module make up with regards to fetal specific eGenes, adult specific eGenes (identified in Gtex v7), or overlapping eGenes (from both fetal and adult datasets). Importantly, co-variates are not driving module clustering. Modules are also not enriched for eGenes, as eGene discovery is a function of sample size in the discovery data set and not biology specific. **(B)** Top 2 Gene Ontology biological process terms enriched for each module. The X axis depicts the −log10(FDR) value, the dotted black line indicates significance (-log_10_(0.05)) with all listed categories significant. **(C)** Top 2 cell types enriched for each module. The X axis depicts the –log10(FDR) value, the dotted black line indicates significance (-log10(0.05)). Cell type markers from human and mouse brain were downloaded from (Hawrylycz et al., 2015; Lein et al., 2007; Mancarci et al., 2017; Miller et al., 2014; Tasic et al., 2016; Winden et al., 2009; Y. Zhang et al., 2014; Y. Zhang et al., 2016) as well as the fetal types from (Polioudakis et al., 2018) **(D-F)** Top hub genes along with edges supported by co-expression are shown for the blue, red, and yellow module. Hub genes are defined by being the top 30 most connected genes based on kME intermodular connectivity. **(G)** Per module QQ plot of SCZ GWAS SNP p-values in regulatory regions defined by eQTLs. The red module and the blue module show the most inflation for enrichment of SCZ GWAS hits. **(H)** Per module QQ plot of ASD GWAS SNP p-values in regulatory regions defined by eQTLs. ASD shows a different pattern than SCZ, with yellow module showing the most inflation for enrichment of ASD GWAS hits. **(I)** Per module rare variant enrichment for SCZ, ASD, ID, Developmental Disorder.

Fetal brain gene co-expression networks, which define core biological processes occurring during cortical development are a useful way to identify processes enriched with common risk variants for neurodevelopmental disorders (Parikshak et al., 2015; Parikshak et al., 2013). However, most risk variants lie within noncoding regions, making their assignment to genes difficult, especially since many regulatory interactions do not involve the closest gene (Won et al., 2016). We used the identified eQTLs to link noncoding variants with target genes and asked whether there were any modules were enriched for ASD and SCZ GWAS signal (Methods). We identify significant enrichments for SCZ (blue p-value = 0.000999) and a marginal trend towards enrichment for ASD (yellow p-value = 0.061) GWAS studies (**Figure 5G and 5H, Figure S8**). Interestingly, the blue module, which shows eQTL/regulatory region GWAS enrichment for SCZ, corresponds to the biological function of neurogenesis as defined by GO analysis, and includes key genes such as DLX1 (essential for GABAergic interneuron production), FGF2 (anteroposterior neural patterning), LHX6 (transcriptional regulator of differentiation and development of neural cells), and SMAD1(progenitor proliferation and differentiation), many of whose enhancer regions have not been previously defined. The yellow module, which shows suggestive eQTL/regulatory region GWAS enrichment for ASD, corresponds to expression regulation defined by GO terms of chromatin organization and mRNA splicing in fetal neurons, and includes key genes such as MEF2A and MEF2D (myocyte enhancer factor 2A and 2D transcription factors both involved in neuronal differentiation), SP2 (transcriptional repressor), HNRNPH3 ( heterogeneous nuclear ribonucleoprotein associated with pre-mRNA processing), and FOXP4 (transcription regulation in brain development). This analysis demonstrates the power of eQTLs, when integrated with co-expression modules, to define where common genetic variation associated with a disease acts through regulation of genes with similar biological functions and potentially similar regulatory control (e.g. transcription factors).

Next, to determine if there was functional overlap in regulatory regions between common GWAS defined annotations above, and rare genetic variation, we examined whether genes harboring rare mutations associated with risk for early onset neurological disease converged on any biological process defined by the fetal co-expression modules (**Figure 5I**). We compiled lists of candidate genes from whole exome sequencing (WES) studies identifying rare and de novo genetic risk variants for ASD, SCZ, intellectual disability, and developmental disorder (Methods; T. N. Turner et al., 2017). Interestingly, we find only the red module enriched for rare variation for SCZ (nominal p-value 0.037), which is also enriched in common variation from SCZ GWAS. The yellow module also exhibits the most significant enrichment for rare variation for ASD (FDR p-value 0.011), also corresponding to the top module enriched in common variation from the ASD GWAS. These analyses provide evidence for overlap in the genes and pathways impacted by common and rare genetic variation in these disorders.

### SCZ TWAS

To further leverage these data to refine and characterize neuropsychiatric disease loci with developmental origins, we used an imputation-based transcription-wide association study (TWAS; (Gusev et al., 2016), to integrate cis-eQTLs and GWAS loci to identify genes whose expression is correlated with schizophrenia. The single previous TWAS for a neurodevelopmental disorder, SCZ, relied on blood, adipose, and adult brain signals, and did not have access to fetal brain eQTL (Gusev et al., 2018). Given the neurodevelopmental origin of these disorders (de la Torre-Ubieta et al., 2018; Gulsuner et al., 2013), we reasoned that these fetal data would provide new perspectives on SCZ risk. We used the summary statistics from Psychiatric Genomics Consortium (PGC) schizophrenia (SCZ) GWAS (Schizophrenia Working Group of the Psychiatric Genomics, 2014) consisting of 79,845 individuals and our fetal brain eQTL data set to identify genes and splicing-events whose imputed cis-regulated expression is associated with SCZ. Given the expression weights, PGC SCZ GWAS Z-scores, and linkage-disequilibrium (LD) reference panel, we computed TWAS statistics (Methods). Of the 2,513 genes and 4,002 introns with significant cis-heritability, we identified 39 gene-SCZ and 58 intron-SCZ (corresponding to 46 unique genes) significant transcriptome-wide associations at Bonferroni corrected p-value <0.05/2,513 for genes and p-value <0.05/4,002 for introns (**Figure 6A, Table S5**). We find 6 genes (YPEL3, SF3B1, NT5DC2, TCTN1, PCDHA2, NDUFA6-AS1) having both a gene-level and intron-level association.

**Figure 6:**
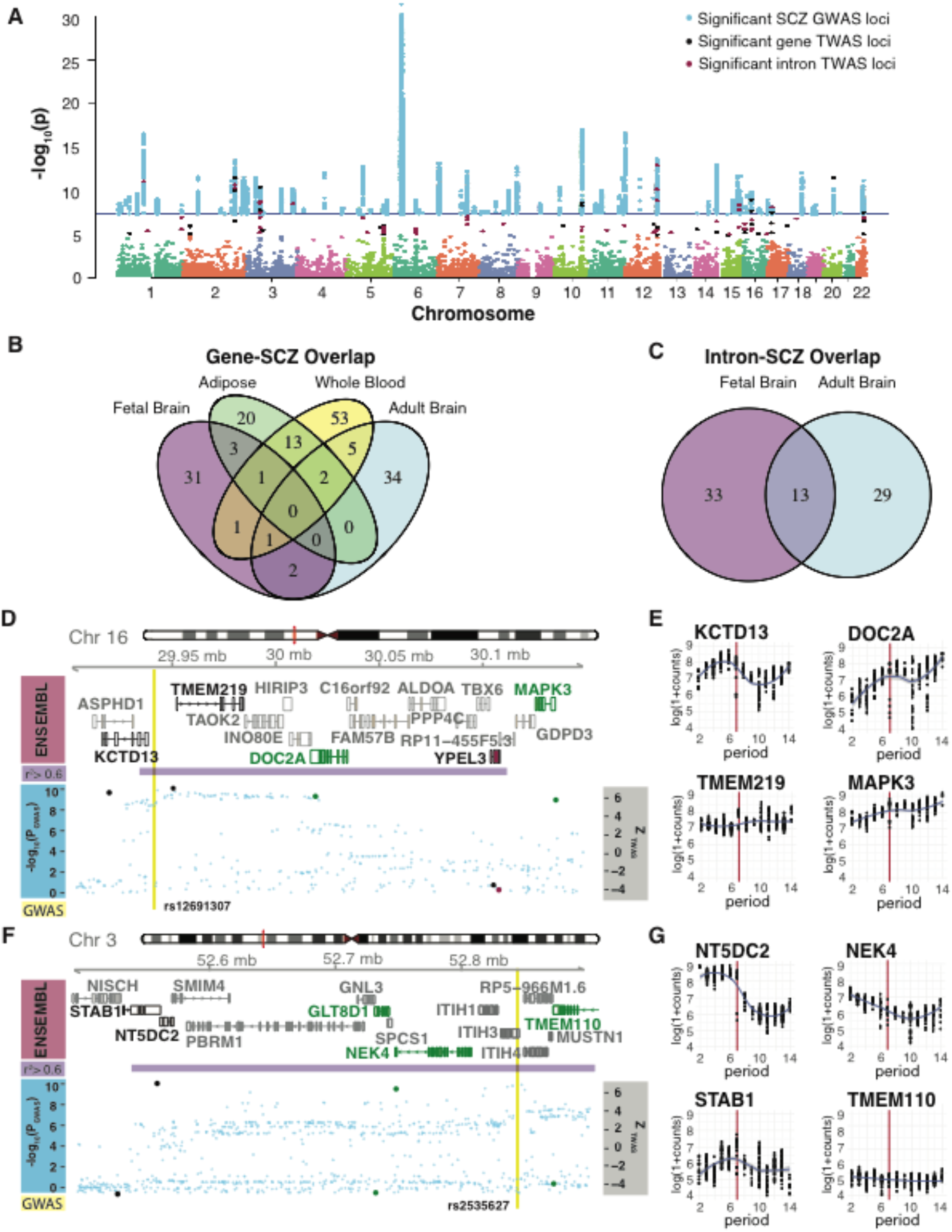
Fetal Brain Schizophrenia TWAS Associationsg. **(A)** Manhattan plot of TWAS results from both genes and intron SCZ associations, highlighting the significant genes (39) and introns (58) at Pbonferroni<0.05, overlaid with only the SCZ PGC GWAS significant hits. Key depicts significant GWAS loci (blue), significant gene level TWAS loci (black), and significant intron level TWAS loci (red). **(B)** Overlap of gene level TWAS hits from fetal brain-SCZ, adult brain-SCZ, adipose-SCZ, and whole blood-SCZ from (Gusev et al., 2018). **(C)** Overlap of intron level TWAS hits from fetal brain-SCZ and adult brain-SCZ from (Gusev et al., 2018). **(D, F)** Illustration of two genomic regions harboring at least one fetal TWAS hit. The Y axis shows Ensemble gene ID, LD, TWAS results overlapped with GWAS hits (-log10(p-value); gene level TWAS hits are black dots, intron level TWAS hits are red dots, and GWAS SNPs are blue dots) and SCZ GWAS hit (yellow line). Ensembl gene names and gene models are colored by significance in datasets; fetal brain gene-SCZ associations (black), fetal brain intron-SCZ associations (red), adult brain gene-SCZ associations (Common Mind Consortium; green). SCZ GWAS hits are shown in yellow, with the LD block surrounding the hit in purple defined by r^2^>0.6. **(E, G)** Corresponding BrainSpan (BrainSpan, 2013) gene expression trajectories throughout human life span of highlighted TWAS genes. In general, these trajectories match expectations in that adult genes, such as MAPK3 and DOC2A have higher adult expression, while NT5DC2 and STAB1 have higher fetal expression and are identified in the fetal analysis, but this is not always the case, as for NEK4 which shows the opposite trajectory. X-axis corresponds to developmental periods defined in (H. J. Kang et al., 2011), where periods 1-7 span embryonic, early fetal, mid-fetal, and late fetal post-conceptional weeks. Periods 8-11 correspond to neonatal and early infancy, late infancy, early childhood, and middle-late childhood. Periods 12-15 correspond to adolescence and adulthood. The red line represents period 8, corresponding to birth.

The previous TWAS study in SCZ referred to above (Gusev et al., 2018), which used the PGZ SCZ GWAS, identified 247 significant transcriptome-wide gene-SCZ and intron-SCZ associations (Gusev et al., 2018). When comparing the significant fetal brain gene-SCZ associations, only 7 genes overlapped between fetal-gene-SCZ and adult-gene-SCZ associations from all three *Gusev. et al* reference panels, with 3 of these 7 genes coming from the adult brain gene-SCZ associations (MAIP1, PCDHA2, and SF3B1) (**Figure 6B**). Thirteen genes with intron-SCZ association overlapped between the 58 fetal brain intron-SCZ associations and the 80 adult brain intron-SCZ associations (**Figure 6C**). The three genes implicated by both fetal and adult TWAS gene associations are protocadherin alpha 2 (PCDHA2), splicing factor 3b subunit 1 (SF3B1) and matrix AAA peptidase interacting protein 1 (MAIP1; C2orf47). PCDHA2 expressed in the brain has been shown to localize to synaptic junctions and play a role in establishing neuronal connections. (Wu & Maniatis, 1999). SF3B1 has been associated with SCZ from the PGC GWAS and has further support from differential expression in a rat psychosis model (Ingason et al., 2015). MAIP1 (C2orf47) plays a role in mitochondrial Ca2+ handling and cell survival and has been associated with SCZ from GWAS performed within a Swedish sample *(N* = 11,244) (Ripke et al., 2013).

A major advantage of TWAS is the identification of genes that did not pass genome-wide significance in the GWAS study. We next examined the overlap between the 108 genome-wide significant GWAS loci (Schizophrenia Working Group of the Psychiatric Genomics, 2014) and significant TWAS associations, in order to identify new risk regions for SCZ. Of the 39 associated genes and 58 associated splice-junctions, 30 genes and 44 splice-junctions (+−500kb) were located within one of the 108 SCZ GWAS published associated regions (Methods), accounting for 27 GWAS region hotspots, with the remaining 9 genes and 14 introns identified by TWAS implicating novel SCZ targets accounting for 17 new risk regions. Of these 17 new risk regions, only 2 were previously identified (+-500k) among the 24 novel loci identified in *Gusev et al.* One of the new SCZ candidate genes that is unique to the fetal dataset is BAIAP2, brain-specific angiogenesis inhibitor 1-associated protein 2 involved in regulation dendritic spine morphogenesis (Choi et al., 2005) and is associated with childhood ADHD and Autism Spectrum Disorder (Liu et al., 2013; Toma et al., 2011). Similar to GWAS studies, TWAS gene-trait association statistics in risk regions are correlated due to LD structure, and thus require fine-mapping for further interpretation. We applied FOCUS (Mancuso et al., 2017) to our fetal-gene TWAS results that lie within a GWAS region to identify a credible set of causal genes at the SCZ GWAS significant loci (Methods) and identified a credible gene set consisting of 20 genes, including KCTD13 and NT5D2. KCTD13 and NT5D2 are both fetal specific hits in GWAS regions where adult specific hits also lie and show greater expression in prenatal time points compared to postnatal in the BrainSpan gene expression trajectories. (**Figure 6D-G, Table S6**, (BrainSpan, 2013)).

We compared the overlap of significant GWAS regions with fetal (39 gene 58 intron) and adult brain (44 gene 80 introns) significant TWAS associations (Methods). At 18 loci there contained at least 1 gene or intron association from both fetal and adult hits. At 9 GWAS regions only adult TWAS gene/introns were implicated, and at 12 GWAS regions only fetal gene/introns were implicated (**Table S7**). For the 18 regions with both adult and fetal implicated gene/introns, 3 of these GWAS loci implicated the same gene, while 8 GWAS loci implicate the same gene that contains a splicing-events, consistent with expectations that using expression panels from different developmental time points or tissues will implicate different genes associated with disease variants (Gusev et al., 2018). It has been shown that tissue of the expression panel used in TWAS studies has a substantial influence on the results (Wainberg et al., 2017), hence the importance of using disease relevant tissues, which for Schizophrenia includes developing brain.

### Intracranial Volume TWAS

Given the importance of cortical neurogenesis, which peaks during mid-gestation (H. J. Kang et al., 2011; Miller et al., 2014), in brain evolution and brain size (Bae, Jayaraman, & Walsh, 2015; Geschwind & Rakic, 2013; Kostovic & Jovanov-Milosevic, 2006; Rakic, 1995, 2009), we reasoned that these eQTL data would be particularly valuable in defining loci involved in intracranial volume in humans. By using fetal brain expression weights, we could specifically highlight early acting genes with a candidate mechanism for which these genetic association signals during time frame crucial to the expansion of the cerebral cortex (de la Torre-Ubieta et al., 2018). We leveraged the GWAS summary statistics of intracranial volume, a meta-analysis of Cohorts for Heart and Aging Research in Genomic Epidemiology (CHARGE) and Enhancing NeuroImaging Genetics through Meta-Analysis (ENIGMA) consortia consisting of 26,577 individuals to perform a transcription-wide association study, as intracranial volume is directly related to brain growth early in brain development (Adams et al., 2016). Of the 2,513 genes and 4,006 introns with significant cis-heritability, we identified 8 genes whose expression (NSF, LRRD1, LRRC37A, LRRC37A2, LRRC37A17P, CENPS, RNF123, AC068152.1) and 7 introns whose splicing is (NT5C2, CRHR1, LRRC37A, LRRC37A2, ARL17A, ARL17B, USP4) significantly associated with intracranial volume at Bonferroni corrected p-value <0.05/2,513 for genes and p-value <0.05/4,006 for introns (**Figure 7A, Table S8**, Methods). An intriguing candidate is NSF, an ATPase involved in vesicular transport and is implicated in having a role in learning, cognition, and memory. It has also been associated with schizophrenia through differential gene expression analysis done in prefrontal cortex and is located in a CNV, also including LRRC37A, LRRC37A2, ARL17A, ARL17B, associated with both dyslexia disorder and Parkinson’s disorder (**Figure 7B**) (Fan et al., 2017; Latourelle, Dumitriu, Hadzi, Beach, & Myers, 2012; Mirnics, Middleton, Marquez, Lewis, & Levitt, 2000; Veerappa, Saldanha, Padakannaya, & Ramachandra, 2014). Another interesting candidate is RNF123, a ubiquitin ligase involved in cell cycle progression found to be overexpressed in the cortex of patients with psychotic depression (Teyssier, Rey, Ragot, Chauvet-Gelinier, & Bonin, 2013).

**Figure 7:**
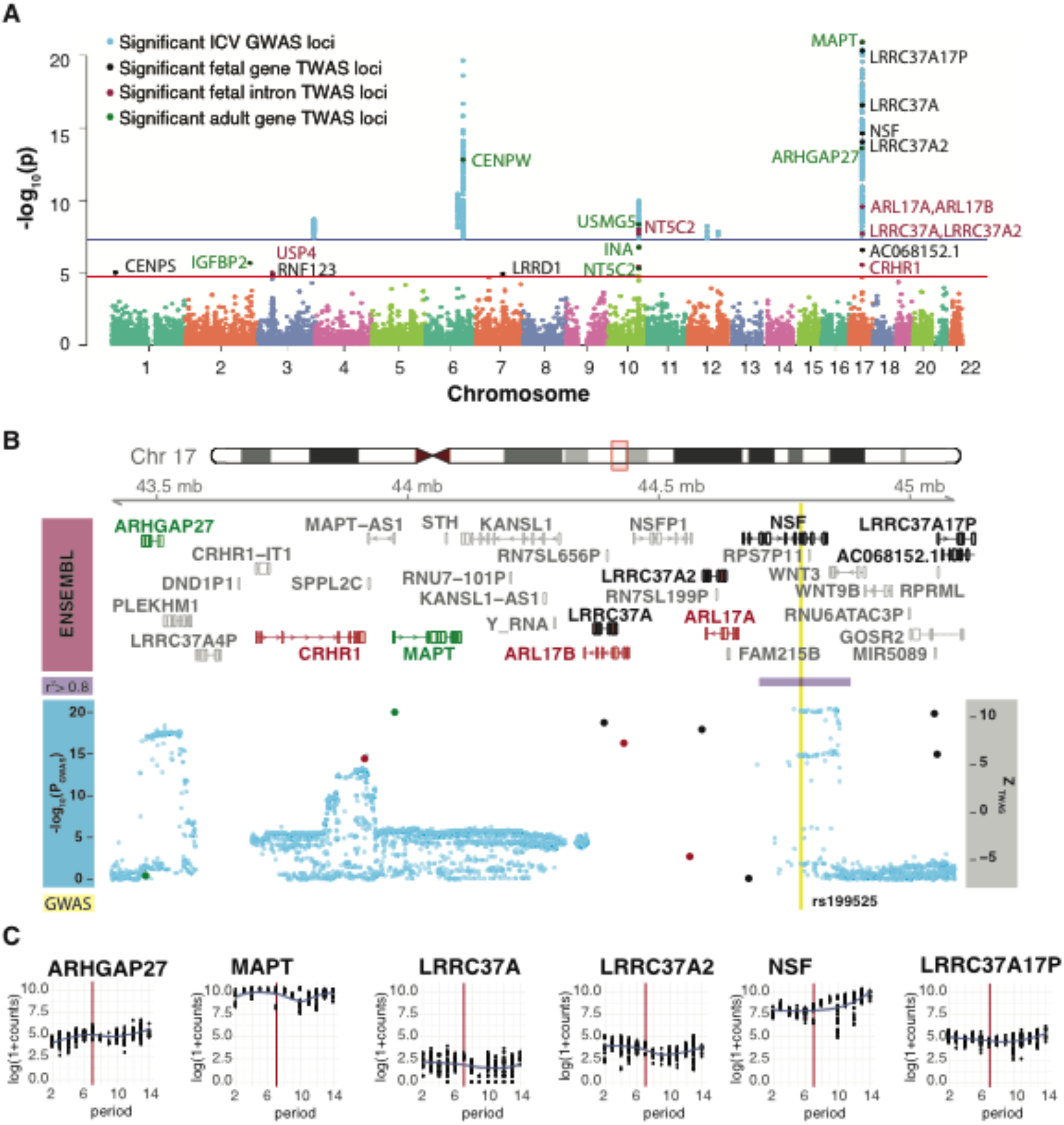
Fetal Brain Intracranial Volume TWAS Associations. **(A)** Manhattan plot of TWAS results from fetal brain gene- and intron-ICV associations and adult brain gene-ICV associations (CommonMind; (Gusev et al., 2016), highlighting the significant fetal brain genes (8) and introns (7) and adult brain genes (7) at Pbonferroni<0.05, overlaid with only the ICV GWAS significant hits. Key depicts significant GWAS loci (blue), significant fetal gene level TWAS loci (black), significant intron level TWAS loci (red), and significant adult gene level TWAS loci in green. **(B)** Illustration of the GWAS locus hotspot on chromosome 17 harboring 5 fetal gene-ICV associations (black), 3 fetal intron-ICV associations (red), adult brain gene-ICV associations (Common Mind Consortium; green). The Y axis shows Ensemble gene ID, LD, TWAS results overlapped with GWAS hits (-log10(p-value); fetal gene level TWAS hits are black dots, adult gene level TWAS hits are green dots, fetal intron level TWAS hits are red dots, and GWAS SNPs are blue dots) and SCZ GWAS hit (yellow line). Ensembl gene names and gene models are colored by significance in datasets; fetal brain gene-SCZ associations (black), fetal brain intron-SCZ associations (red), adult brain gene-SCZ associations (Common Mind Consortium; green). The ICV GWAS hit is shown in yellow with the LD block surrounding the hit in purple defined by r^2^>0.8. **(C)** Corresponding BrainSpan (BrainSpan, 2013) gene expression trajectories throughout human life span of highlighted TWAS genes. X-axis corresponds to developmental periods defined in (H. J. Kang et al., 2011), where periods 1-7 span embryonic, early fetal, mid-fetal, and late fetal post-conceptional weeks. Periods 8-11 correspond to neonatal and early infancy, late infancy, early childhood, and middle-late childhood. Periods 12-15 correspond to adolescence and adulthood. The red line represents period 8, corresponding to birth.

We also ran the TWAS using the published adult brain weights calculated from the Common Mind Consortium (Gusev et al., 2018).Of the 5,376 genes with significant cis-heritability, 7 genes (NT5C2, INA, USMG5, ARHGAP27, MAPT, IGFBP2, CENPW) are significantly associated with intracranial volume (Bonferroni corrected p-value <0.05/5,376), none of which overlap the fetal identified genes, except for NT5C2 at the fetal intron level (**Figure 7A**). NT5C2 is a hydrolase that plays a role in cellular purine metabolism that has been linked to intellectual disability and spastic paraplegia (Darvish et al., 2017).

## Discussion

Our analysis provides the first genome wide map of human eQTLs and spliceQTLs during cerebral corticogenesis, a critical epoch of brain development, expanding our understanding of complex gene and splicing regulation in the developing brain. We have comprehensively described the implicated regulatory regions defined by eSNPs and sSNPs, enabling the integration of developmental diversity to previous adult brain functional genomic analysis and genetic variant interpretation. We show that much of the genetic variation controlling regulation of expression and splicing in the human brain is sensitive to two distinct periods of developmental stage. In the context of early onset neurological and psychiatric disorders, this provides a new window into genetic control of gene expression and splicing regulation during a critical time point for disease development.

We find that integrating eQTLs through regulatory region annotation with WGCNA defined modules from fetal brain expression shows GWAS disease enrichment for both ASD and SCZ among specific biological processes, indicating specific convergent biology. This substantially advances previous work based on rare, gene disrupting de novo mutations (Iossifov et al., 2012; Ruzzo et al., 2018; Sanders et al., 2012) by showing convergence in genetic risk factors, even at the level of common variants that lie in regulatory regions. We show genes involved in chromatin organization and splicing are enriched for ASD GWAS loci in their eQTL defined regulatory regions, similar to rare variants. For SCZ, we find neurogenesis and central nervous system development enrichment, parallel to fetal brain Hi-C analysis (Won et al., 2016), providing independent support for neurogenesis as a key process in SCZ risk. Furthermore, we show that SCZ GWAS enrichment from partitioned heritability significantly differs when using fetal eQTL defined genome annotations versus adult eQTL defined genome annotations, consistently indicating the fetal annotations to be the highest enriched among all functional annotations. Similar significant enrichment of SCZ partitioned heritability has been shown for fetal brain ATAC-seq peaks (de la Torre-Ubieta et al., 2018), demonstrating the multiple lines of evidence that regulatory regions active in early cortical development harbor significant SCZ risk and are likely to be a crucial site of action for disease risk.

Finally, by integrating our map of fetal gene expression and splicing regulation with the SCZ and intracranial volume GWAS studies through application of TWAS, we are able to identify new candidate genes and candidate molecular mechanisms through which these disease associated common variants may be acting. We discover several genes that lie in significant GWAS loci, but additionally, we discover new regions of the genome associated with SCZ through aggregation of GWAS signal among from the significant genetic cis-predictors of genes expression. Moreover, when comparing the implicated genes and introns from fetal brain expression weights with genes and introns implicated via adult brain expression weights, there is very little overlap. To some degree this is expected and reflects TWAS’s known sensitivity to the specific expression panel, as well as the differences in eGenes and effect sizes between fetal and adult eQTLs shown in our analysis. It does, however, highlight the importance of picking appropriate expression studies for a given phenotype. Here, the SCZ risk genes predicted by TWAS based on fetal brain expression contributes to the growing list of candidate genes discovered from adult expression for a more complete view of potentially casual genes and splicing events. These results and others (de la Torre-Ubieta et al., 2018; Won et al., 2016) are very consistent with the seminal framing of SCZ as a neurodevelopmental disorder (Weinberger, 1987).

In this regard, GWAS regions that yield discordant significant TWAS gene associations depending on the developmental time period of reference panel are interesting to consider. A particularly salient example is the region containing the 16p11.2 copy number variant associated with multiple neurodevelopmental conditions such as SCZ, ASD, ID, and BIP, (Bernier et al., 2017; Hanson et al., 2015; Shinawi et al., 2010; Weiss et al., 2008; W. Zhou et al., 2018). Here, we find KCTD13, TMEM219, and YPEL3 highlighted by fetal brain expression with fine mapping of this loci implicating KCTD13, MVP, and AC120114.1, whereas MAPK3 and DOC2A, both of which are also in the copy variant region, are implicated by adult brain expression. While this CNV region contains 29 genes, KCTD13 has been studied extensively due and has been posited as a major drivers to phenotypic changes based on experimental manipulation (Golzio et al., 2012; Luo et al., 2012), but the relationship is unlikely to be simple or involve only one gene (Escamilla et al., 2017; Golzio et al., 2012). While more work is necessary to understand the role of KCTD13, MAPK3, and the 16p11.2 CNV, our TWAS results suggest these genes may play a role in disease biology through action at different developmental time periods. These results further demonstrate the importance of considering developmental stage of brain expression and regulation when using eQTL data to interpret disease associated variants. They also highlight the value of using these data to further characterize developmental time points during which genetic disruption acts for neurodevelopmental and early onset neuropsychiatric diseases, suggesting that more detailed maps of the effects of common genetic variation effects throughout the lifespan will be of value.

## Acknowledgements

Fetal tissue was collected from the UCLA CFAR (5P30 AI028697). RNA-seq libraries were prepared and sequenced by the UCLA Neuroscience Genomics Core. We thank members of the Geschwind lab for helpful discussions and critical reading of the manuscript. This work was supported by the NIH (5R37 MH060233, 5R01 MH094714, 1R01 MH110927 to D.H.G, R00MH102357 to J.L.S, 1F32MH114620 to G.R) and the National Institute of Neurological Disorders And Stroke of the National Institutes of Health Training Grant (T32NS048004 R.L.W.).

## Author Contributions

L.T.-U., J.L.S., and D.H.G. designed the study. D.H.G. oversaw all of the analyses and supervised the study. L.T.-U. and J.L.S. performed sample collection and brain dissections, genotyping and RNA-seq library preparation. R.L.W. processed and managed the data, performed eQTL analysis, interpreted results, and performed integration with other expression and epigenetic data, GWAS, and network analysis, with input from G.R., C.H., M.J.G., N.M. and B.P. N.M. performed TWAS fine mapping. R.L.W. and D.H.G. wrote and prepared the manuscript, with feedback from all authors.

## Tables

<u>Table S1. Related to Figure 2:</u> Statistics for significant eQTLs, top SNP per gene. *gene_id* and *hgnc_symbol* is the Ensembl gene ID from GRCh37 and corresponding gene symbol. *gene_chr, gene_start, gene_end* are the genomic locations of the Ensembl gene ID from GRCh37 and hg19. *num_var* is the number of variants tested in the cis-window. *beta_shape1* and *beta_shape2* are the MLE of the shape 1 and shape 2 parameter of the Beta distribution. *variant_id* is the best SNP (smallest p-value) in the form chr_position_ref_alt in hg19 corrdinates. *tss_distance* is the distance between the variant and the TSS of the gene. *Chr, pos, ref, alt* correspond to the variant in hg19 coordinates. *pval-nominal* is the nominal p-value of the association. *slope* is the regression coefficient associated with the nominal p-value association. *pval_beta* is the permutation p-value obtained via beta approximation. *qval* is the Storey & Tibshirani corrected permutation p-value.

<u>Table S2. Related to Figure 3</u>: Statistics for significant sQTLs, top SNP per intron. *intron_id* is the Leafcutter intron identification in the form chr:intron_start:intron:end:cluster_id. *hgnc_symbol* is the gene symbol of the gene the intron maps to, since Leafcutter is annotation free intron clusters may map to multiple genes if there is overlap. *intron_chr, intron_start, intron_end* are the genomic locations of the intron in hg19 coordinates. *num_var* is the number of variants tested in the cis-window. *beta_shape1* and *beta_shape2* are the MLE of the shape 1 and shape 2 parameter of the Beta distribution. *variant_id* is the best SNP (smallest p-value) in the form chr_position_ref_alt in hg19 corrdinates. *tss_distance* is the distance between the variant and the intron start. *Chr, pos, ref, alt* correspond to the variant in hg19 coordinates. *pval-nominal* is the nominal p-value of the association. *slope* is the regression coefficient associated with the nominal p-value association. *pval_beta* is the permutation p-value obtained via beta approximation. *qval* is the Storey & Tibshirani corrected permutation p-value.

<u>Table S3. Related to Figure 3</u>: 47 genes with the same top SNP both their eQTL and sQTL. ENSID and Gene is the Ensembl gene ID from GRCh37 and corresponding gene symbol. SNP is the top SNP for the eQTL and sQTL. Sqtl_pval and eqtl_pval are the corresponding FDR corrected p-vaules (qvalue).

<u>Table S4. Related to Figure 5</u>: WGCNA Module Membership. *ENSG* and *Gene* is the Ensembl gene ID from GRCh37 and corresponding gene symbol. *Gene.colors* is the color of the module in which the gene belongs. The next 18 columns are the module kME of the corresponding module color. *eQTL Pvalue* is the nominal p-value for the top eQTL if that gene contains a significant eQTL.

<u>Table S5. Related to Figure 6</u>: SCZ TWAS gene and intron hits. *Gene* is the gene symbol of the significant TWAS hit. *Chr, gene/intron start, gene/intron end* correspond to the gene or intron’s postion in hg19 coordinates. *BEST.GWAS.ID* and *BEST.GWAS.Z* are the most significant SCZ GWAS SNP in the locus and correspond Z-score of the SCZ GWAS SNP. *TWAS.Z* and *TWAS.P* are the TWAS Z-score and p-value. *Expression* designates which expression weights were used, gene or intron level.

<u>Table S6. Related to Figure 6</u>: SCZ Focus fine mapping of SCZ TWAS hits overlapping PGC SCZ GWAS significant regions. *Gene* and *ID* are the gene symbol and corresponding Ensembl gene ID from GRCh37. *BLOCK* is the genomic window in the form chr:region_start.region_stop corresponding to LD blocks around the PGC SCZ GWAS hits. *RESID.Z* is the TWAS Z score after accounting for the estimated intercept at each region. *PIP* is the posterior inclusion probability (causality probability). *IN.CRED.SET* marks if a gene or null model are included in the 90% credible gene set. If a gene set contains the null model in the 90% credible gene set it is not included.

<u>Table S7. Related to Figure 6</u>: TWAS hits overlapping GWAS regions from fetal and adult expression panels. Ensid and gene is the Ensembl gene ID and corresponding gene symbol. GWASsnp corresponds to one of the 108 PGC SCZ GWAS loci. Annotation indicates which expression panel the TWAS hit is coming from (fetal vs adult) and what type of expression (gene-level or splicing). At 18 loci there contained at least 1 gene or intron association from both fetal and adult hits. At 9 GWAS regions only adult TWAS gene/introns were implicated, and at 12 GWAS regions only fetal gene/introns were implicated.

<u>Table S8. Related to Figure 7</u>: ICV TWAS gene and intron hits. *Gene* is the gene symbol of the significant TWAS hit. *Chr, gene/intron start, gene/intron end* correspond to the gene or intron’s postion in hg19 coordinates. *BEST.GWAS.ID* and *BEST.GWAS.Z* are the most significant ICV GWAS SNP in the locus and correspond Z-score of the ICV GWAS SNP. *TWAS.Z* and *TWAS.P* are the TWAS Z-score and p-value. *Expression* designates which expression weights were used, gene or intron level.

## STAR Methods

Key Resources Table
| REAGENT or<br>RESOURCE | SOURCE | IDENTIFIER |
| --- | --- | --- |
| **Critical** |  |  |
| **Commercial** |  |  |
| **Assays** |  |  |
| miRNA easy-mini | QIAGEN |  |
| DNeasy Blood and<br>Tissue Kit |  |  |
| Stranded TruSeq<br>kit with Ribozero<br>Gold | Illumina |  |
| **Deposited Data** |  |  |
| Raw and analyzed<br>data: RNA-seq | This paper |  |
| Raw and analyzed<br>data: DNA | This paper |  |
| Hg 19 genome<br>build | N/A | ftp://ftp-trace.ncbi.nih.gov/1000genomes/ftp/technical/refere<br>nce/phase2_reference_assembly_sequence/ |
| Roadmap Epigenomics 25-state model | (Roadmap Epigenomics et al., 2015) | https://personal.broadinstitute.org/jernst/MODEL_IMPUTED12MARKS/ |
| Gencode v 19 | (Harrow et al., 2012) | https://www.gencodegenes.org/releases/19.html |
| ENSEMBL Human genome assembly | N/A | ftp://ftp.ensembl.org/pub/release-75/fasta/homo_sapiens/dna/ |
| Developing human cortex Hi-C | (Won et al., 2016) | https://www.ncbi.nlm.nih.gov/geo/query/acc.cgi?acc=GSE77565 |
| Developing human cortex ATAC-seq | (de la Torre-Ubieta et al., 2018) | https://doi.org/10.1016/j.cell.2017.12.014 |
| GTEx v7 | (Consortium et al., 2017) | https://www.gtexportal.org/home/datasets<br>https://www.ncbi.nlm.nih.gov/projects/gap/cgi-bin/study.cgi?study_id=phs000424.v7.p2 |
| HapMap3 | (International HapMap, 2003) | ftp://ftp.ncbi.nlm.nih.gov/hapmap/ |
| 1000 Genomes Project Phase3 | (Genomes Project et al., 2015) | ftp://ftp.1000genomes.ebi.ac.uk/vol1/ftp/release/20130502/ |
| BrainSpan | (BrainSpan, 2013) | http://brainspan.org/ |
| Transcription factor binding sites | (Arbiza et al., 2013) |  |
| RNA binding<br>protein binding<br>sites | (Y. C. Yang et al., 2015) | https://omictools.com/clipdb-tool |
| Functional Gene<br>Constraint Scores | (Lek et al., 2016) | http://exac.broadinstitute.org |
| **Software and Algorithms** |  |  |
| R (v3.2.1) |  | https://www.r-project.org/ |
| FastQC | N/A | https://www.bioinformatics.babraham.ac.uk/projects/fastqc/ |
| RNA STAR v2.4 | (Dobin et al., 2013) | https://github.com/alexdobin/STAR |
| Picard Tools<br>v1.139 | N/A | http://broadinstitute.github.io/picard/ |
| Samtools v1.2 | (H. Li et al., 2009) | https://samtools.github.io |
| PLINK 1.9 | (Chang et al., 2015) | https://www.cog-genomics.org/plink2 |
| HTseq Counts | (Anders, Pyl, & Huber, 2015) | https://htseq.readthedocs.io/en/release_0.10.0/ |
| VerifyBamID<br>v1.1.2 | (Jun et al., 2012) | https://genome.sph.umich.edu/wiki/VerifyBamID |
| Plinkseq v0.10 | N/A | https://www.atgu.mgh.harvard.edu/plinkseq |
| ComBat | (W. E. Johnson, Li, & Rabinovic, 2007) | https://www.bu.edu/jlab/wp-assets/ComBat/Abstract.html |
| WGCNA | (Langfelder & Horvath, 2008; B. Zhang & Horvath, 2005) | https://cran.r-project.org/web/packages/WGCNA/index.html |
| Beagle v4.1 | (Browning & Browning, 2016) | https://faculty.washington.edu/browning/beagle/beagle.html |
| GATK v3.5 | (Van der Auwera et al., 2013) | https://software.broadinstitute.org/gatk/ |
| HCP | (Mostafavi et al., 2013) | https://github.com/mvaniterson/Rhcpp |
| FastQTL | (Ongen et al., 2016) | http://fastqtl.sourceforge.net |
| Matrix eQTL | (Shabalin, 2012) | https://cran.r-project.org/web/packages/MatrixEQTL/index.html |
| EMMAX | (H. M. Kang et al., 2008) | http://genetics.cs.ucla.edu/emmax_jemdoc/ |
| METAL<br>v3.25.2011 | (Willer et al., 2010) | http://csg.sph.umich.edu/abecasis/Metal/ |
| Leafcutter | (Y. I. Li et al., 2018) | http://davidaknowles.github.io/leafcutter/index.html |
| GREGOR | (Schmidt et al., 2015) | https://genome.sph.umich.edu/wiki/GREGOR |
| Variant Effect<br>Predictor | (McLaren et al., 2016) | https://uswest.ensembl.org/info/docs/tools/vep/index.html |
| LD Score Regression | (Finucane et al., 2015) | https://github.com/bulik/ldsc |
| FUSION | (Gusev et al., 2016) | http://gusevlab.org/projects/fusion/ |
| FOCUS | (Mancuso et al., 2017) | http://github.com/bogdanlab/focus/ |
| GenomicRanges Package v1.16.4 | (Lawrence et al., 2013) | http://bioconductor.org/packages/release/bioc/html/GenomicRanges.html |
| Gviz Package v1.8.4 | (Hahne & Ivanek, 2016) | http://bioconductor.org/packages/release/bioc/html/Gviz.html |
| biomaRt Package v2.20.0 | (Durinck, Spellman, Birney, & Huber, 2009) | http://bioconductor.org/packages/release/bioc/html/biomaRt.html |
| BSgenome.Hsapiens.UCSC.hg19 v1.32.0 | N/A | http://bioconductor.org/packages/release/data/annotation/html/BSgenome.Hsapiens.UCSC.hg19.html |
| qvalue Package | (Storey & Tibshirani, 2003) | https://www.bioconductor.org/packages/devel/bioc/html/qvalue.html |
| Cqn Package v1.26.0 | (Hansen, Irizarry, & Wu, 2012) | https://bioconductor.org/packages/release/bioc/html/cqn.html |
| Dplyr Package v0.7.6 | N/A | https://cran.rstudio.com/web/packages/dplyr/index.html |
| Qqman Package v0.1.4 | (S. D. Turner, 2014) | https://cran.r-project.org/web/packages/qqman/ |
| AnnotationDbi<br>Package v1.42.1 | (Pagès, Carlson,<br>Falcon, & Li,<br>2018) | https://www.bioconductor.org/packages/release/bioc/html/AnnotationDbi.html |
| flashClust Package<br>v1.01-2 | (Langfelder &<br>Horvath, 2012) | https://cran.r-project.org/web/packages/flashClust/index.html |
| GenomicFeatures<br>Package v1.42.1 | (Lawrence et al.,<br>2013) | https://bioconductor.org/packages/release/bioc/html/GenomicFeatures.html |
| Igraph Package<br>v1.2.1 | (Nepusz &<br>Csárdi, 2006) | https://cran.r-project.org/web/packages/igraph/index.html |
| Corrplot v0.84<br>Package | (Wei & Simko,<br>2017) | https://cran.r-project.org/web/packages/corrplot/index.html |

## Contact for Reagent and Resource Sharing

Further information and requests for resources and reagents should be directed to and will be fulfilled by the Lead Contact, Daniel H. Geschwind.

## Experimental Model and Subject Details

### Developing Human Brain Samples

Fetal tissue was obtained from the UCLA Gene and Cell Therapy core according to IRB guidelines from 233 donors (post-conception weeks: 14-21) following voluntary termination of pregnancy. This study was performed under the auspices of the UCLA Office of Human Research Protection, which determined that it was exempt because samples are anonymous pathological specimens. Full informed consent was obtained from all of the parent donors.

## Method Details

### Data Generation

Total RNA and genomic DNA from human fetal brain tissue from PCW 14-21 that visually appeared to be cortical was extracted using miRNeasy-mini (Qiagen) and DNeasy Blood and Tissue Kit (DNA) or were extracted using trizol with glycogen followed by column purification. Library preparation via Illumina Stranded TruSeq kit with Ribozero Gold ribosomal RNA depletion library prep was followed by sequencing on 233 brains and genotype array data on 212 brains was generated at the UCLA Neurogenomics Core. Pseudo-randomization to decrease correlation between sequencing lane and biological variables like sex and gestation week were performed. RNA samples were pooled, randomized, and run on 4 lanes. Ribozero, ribosome depleted, 50 bp paired-end RNA sequencing was performed with mean sequencing depth of 60 million reads on an Illumina HiSeq2500.

Genotyping was performed at the UCLA Neurogenomics Core (UNGC) on either Illumina HumanOmni2.5 or HumanOmni2.5Exome platform in 8 batches. SNP genotypes were exported into PLINK format. Batches were merged and markers that did not overlap genotyping platforms were removed. SNP marker names were converted from Illumina KGP IDs to rsIDs using the conversion file provided by Illumina. Quality control was performed in PLINK v1.9 (Chang et al., 2015). SNPs were filtered based on Hardy-Weinberg equilibrium ( --hwe 1e6), minor allele frequency ( --maf 0.01), individual missing genotype rate (--mind 0.10), variant missing genotype rate (--geno 0.05) resulting in 1,799,583 variants (**Figure S1**).

### RNA-sequencing Data Processing Pipeline

All raw RNAseq fastq files, 4 per sample run on different lanes, were run through FastQC (https://www.bioinformatics.babraham.ac.uk/projects/fastqc/). FastQC output was visually inspected and sequencing lanes where the “per tile sequence quality” was red were removed, there was no sample with more than one sequencing lane removed. Fastq files were aligned to the GRCH37.p13 (hg19; Homo_sapiens.GRCh37.75.dna.primary_assembly.fa: ftp://ftp.ensembl.org/pub/release-75/fasta/homo_sapiens/dna/) reference genome using STAR v2.4 (Dobin et al., 2013). SAM files were sorted, indexed, converted to BAM files and merged across lanes from the same sample using Samtools v1.2 (H. Li et al., 2009). Gene quantifications were calculated using HTSeq-counts v0.6.0 (Anders et al., 2015) using an exon union model on the basis of Gencode v19 comprehensive gene annotations (Harrow et al., 2012). Quality control metrics were calculated using PicardTools v1.139 (http://broadinstitute.github.io/picard) and Samtools. A sex incompatibility check was also performed using XIST expression and Y chromosome non-pseudoautosomal expression which are known to show different patterns of expression in males and females. A scatter plot of XIST expression versus the sum of expression of genes in the non-pseudoautosomal region of the Y chromosome showed no gender mismatches. (**Figure S1**)

### Sample Swap Identification

QC’d genotypes and sample BAM files were used to identify any sample identity swaps between the RNA and DNA experiments using VerifyBamID v1.1.2 (Jun et al., 2012). We identified 4 samples where [CHIPMIX] ~ 1 AND [FREEMIX] ~ 0, indicative of unmatching RNA and DNA, which were removed in the VCF file.

### Genotyping Pipeline

PLINK genotype files were converted to vcf files using Plinkseq v0.10 (https://www.atgu.mgh.harvard.edu/plinkseq). Genotypes were imputed into the 1000 Genomes Project phase 3 multi-ethnic reference panel (Genomes Project et al., 2015) by chromosome using Beagle v4.1 (Browning & Browning, 2016) and subsequently merged. Multiallelic sites were removed using GATK v3.5 (Van der Auwera et al., 2013). Imputed genotypes were filtered for Hardy-Weinberg equilibrium p-value < 1 x 10-6 and minor allele frequency (MAF) 5%. Imputation quality was assessed filtering variants where allelic R-squared >0.5 and dosage R-squared >0.5 by GATK, resulting in ~6.6 million autosomal SNPs. We restricted to only autosomal due to sex chromosome dosage, as commonly done (Consortium, 2015).

### RNA-seq Quality Control and Normalization

Gene counts were compiled from HTSeq Count (Anders et al., 2015) quantifications and imported into R version 3.2.1 for downstream analyses. Gene counts were put through quality control, removing genes that were not expressed in 80% of samples with 10 counts or more. Expression was then corrected for GC content, gene length, and quantile normalized to a standard normal distribution, as commonly done in QTL studies (Battle et al., 2015). Sample outliers were removed based on standardized sample network connectivity Z scores < 2 (B. Zhang & Horvath, 2005) (**Figure S2F**). ComBat batch correction was performed (W. E. Johnson et al., 2007). After quality control and normalization, there remained 201 samples with 15,930 genes expressed (on the basis of Gencode v19 annotations) at sufficient levels.

### Covariate Selection

To evaluate and remove global effects of gene expression, we used PLINK v1.9 (Chang et al., 2015) to run multidimensional scaling on the QC’d imputed genotypes and to verify ancestral backgrounds of the samples. We aggregated the final 201 samples with HapMap3 (International HapMap, 2003) of 1397 samples across 11 populations (87 ASW, 165 CEU, 137 CHB, 109 CHD, 101 GIH, 113 JPT, 110 LWK, 86 MXL, 184 MKK, TSI 102, YRI 203). A plot of the first two MDs components of the merged data shows the genetic ancestry of our samples among a diverse reference population (**Figure S2H**). For eQTL analysis, the top 3 MDS components calculated only in the fetal brain samples were used as covariates.

We also looked at the correlation of known measured biological covariates, measured technical covariates, as well as RNA quality control metrics from Picard tools (gestation week, RIN, sex, purification method, 260:230 ratio, 260:280 ratio, read depth, percent chimeras, 5’ bias, 3’ bias, AT dropout) with the top 10 principle components of the expression data and find the top principal component corresponds to the age of the sample (gestation week) and the second component corresponds to the RNA integrity number (RIN) (**Figure S2G**).

To measure hidden batch effects and confounders, hidden covariate analysis was performed using Hidden Covariates with a Prior (HCP) (Mostafavi et al., 2013). Hidden factors were calculated given the known measured factors. HCP was run separately for varying number of inferred hidden components: 5, 10, 15, 20, 25, 30. We included 20 HCPs in our eQTL model which we found to maximized eGene discovery along with gestation week, RIN, and sex (**Figure S3A**). We correlated the 20 HCPs along with gestation week, RIN, sex, and top 3 genotype PCs (all covariates used in the final model) with the measured factors and Picard metrics, as well as the top 20 PCs of expression to gain insight to meaning of the HCPs. We see each inferred hidden component’s relationship to the known variables is complex and distributed across variables.

### cis-eQTL mapping

We performed cis-eQTL mapping using FastQTL (Ongen et al., 2016), a defined cis window of 1 megabase up- and down-stream of the transcription start site for 15,930 expressed genes, and correction for the following covariates: gestation week, RIN, sex, 20 HCPs, 3 genotype PCs. FastQTL (Ongen et al., 2016) was run in the permutation pass mode (1000 permutations) to identify the best nominal associated SNP per phenotype and with a beta approximation to model the permutation outcome (**Figure S4A**) and correct for all SNPs in LD with the top SNP per phenotype. Beta approximated permutation p-values were then multiple test corrected using the q-value Storey and Tibshirani FDR correction (Storey & Tibshirani, 2003). We define eQTL containing genes (eGenes) by having an FDR q-value <=0.05. Secondary, independent eQTLs were identified by rerunning permutation tests in FastQTL for every eGene conditioning on the primary eSNP.

To look for inflation in a Q-Q plot, we randomly chose 10 genes to run eQTL analysis in trans, testing all SNPs genome-wide with association with gene expression using MatrixEQTL (Shabalin, 2012). We corrected for the same covariates as in the cis-eQTL analysis. The Q-Q plot of the trans-eQTLs shows no inflation, an indication that our eQTL results are not confounded by population stratification. (**Figure S4B**)

### EMMAX

To further check that our eQTLs are not due to the population differences of our samples, we ran cis-eQTL analysis with EMMAX (H. M. Kang et al., 2008), which accounts for population structure using a genetic relationship matrix. We used the emmax-kin function (-v -h -s -d 10) to create the IBS kinship matrix. EMMAX was run for each gene with a cis-window of +-1MB around the TSS, correcting for the same covariates in the FastQTL analysis. Nominal EMMAX p-values were corrected for multiple testing using the q-value Storey and Tibshirani FDR correction (Storey & Tibshirani, 2003).

To assess overlap between FastQTL and EMMAX, FastQTL was also run in the nominal pass mode to obtain nominal p-values for all cis-SNPs tested per gene. FastQTL nominal p-values were also corrected for multiple testing using the q-value Storey and Tibshirani FDR correction. We discover 920,356 nominal eQTLs at a 5% FDR threshold. To compare nominal results, we defined eGenes as a gene containing a significant SNP association at FDR <=0.05. We found 93.8% of eGenes from the FastQTL analysis was an eGene in the EMMAX analysis. Additionally, we compared all SNP-gene pairs tested and found 92.8% of significant SNP-gene associations from nominal FastQTL to be significant in the EMMAX analysis (**Figure S5E-F**).

### Meta-Analysis

We split up our sample into six groups based on hierarchical clustering of the top 3 PCs of the genotype data. (Supplemental) The size of each group ranged from 12 samples to 47 samples and each group corresponded to distinct ancestries based on the MDS plots of samples merged with HapMap3. We performed association testing between the top SNP per gene, identified by FastQTL permutation pass eQTL analysis, within group using the lm() function in R, correcting for gestation week, RIN, age, and 20HCPs. A fixed effect meta-analysis was then run between groups using METAL v3.25.2011 (Willer et al., 2010), which implements a Cochran’s Q test for heterogeneity. We find significant heterogeneity at 10% of our eQTLs and find 87% of our eQTLs are significant in the meta-analysis at FDR 0.05% (q-value) strongly suggesting our results are not due to population stratification (**Figure S5B-D**).

### Intron Cluster Quantifications

We used Leafcutter (Y. I. Li et al., 2018) to leverage information from reads that span introns to quantify clusters of variably spliced introns. From the already aligned FASTQ files by STAR, output bam files were converted into junction files. Intron clustering was performed using default settings of 50 reads per cluster and a maximum intron length of 500kb. Clusters then went through quality control consisting of removal of clusters having zero reads in more than 10 individuals, clusters having more than 20 reads in less than 100 individuals, and introns with less than 5 individuals having non-zero counts. The Leafcutter prepare_genotype_table script was then used to calculate intron excision ratios and to filter out introns used in less than 40% of individuals with almost no variation. Intron excision ratios were then standardized and quantile normalized.

### sQTL mapping

Standardized and normalized intron excision ratios calculated by leafcutter was used as the phenotype for sQTL mapping. FastQTL (Ongen et al., 2016) was used to test for association between SNPs within a cis-region of +-100kb of the intron cluster and intron ratios within cluster. Hidden covariate analysis was performed using Hidden Covariates with a Prior (HCP) (Mostafavi et al., 2013) on intron excision ratios given the same known covariates used for eQTL HCP calculations. We included 5 HCPs in our spliceQTL model which we found to maximized intron QTL (**Figure S3C**) discovery along with gestation week, RIN, and sex. FastQTL was run in the permutation pass mode (1000 permutations). Beta approximated permutation p-values were then multiple test corrected using the q-value Storey and Tibshirani FDR correction. We define sQTL as an intron having an FDR q-value <=0.05, and an sGene as a gene containing a significant sQTL at any intron.

### ATAC-seq Overlap of eQTLs

Fetal brain ATAC-seq peaks were obtained from (de la Torre-Ubieta et al., 2018). We annotated eQTLs as being supported by ATAC if the LD block (r2 >0.8 PLINK) around its eSNP overlapped an open chromatin region. To test for significance, we created a null set of eQTLs (q-value > 0.2), annotated overlap with ATAC peaks, and then ran a Fisher’s exact test.

### Hi-C Overlap of eQTLs

Fetal brain CP and GZ Hi-C topological association domain bed files were obtained from (Won et al., 2016). eSNPs located within 10kb of the eGene TSS were removed, as Hi-C cannot detect any chromosomal interaction less than 10kb apart. We defined any remaining eQTL as overlapping Hi-C if the LD block (r2 >0.8 PLINK) around its eSNP fell in one 10kb TAD bins and the corresponding eGene +-2kb overlapped with the other 10kb TAD bin in either CP or GZ. To test for significance, we created a null set of eQTLs (q-value > 0.2), annotated overlap with Hi-C, and then ran a Fisher’s exact test.

### Functional enrichment of QTLs in epigenetic marks, transcription factor, and splicing factor binding sites

We performed functional enrichment of both eQTLs and sQTLs using GREGOR (Genomic Regulatory Elements and Gwas Overlap algoRithm) (Schmidt et al., 2015) to evaluate enrichment of variants in genome wide annotations. We downloaded the 25 state ChromHMM model BED files from the Roadmap Epigenetics Project (Ernst & Kellis, 2015; Roadmap Epigenomics et al., 2015), generated from a set of 5 core chromatin marks assayed in fetal brain. We downloaded consensus transcription factor and DNA-binding protein binding site BED files (Arbiza et al., 2013), which called consensus binding sites from multiple cell types from Encode CHIP-seq data which was used to computationally annotate all possible genome-wide sites for 78 binding proteins. We filtered to 62 binding proteins that showed cortical brain expression in BrainSpan (BrainSpan, 2013). Lastly, we obtained human RNA binding protein (RBP) binding site BED files from CLIPdb (Y. C. Yang et al., 2015) database of publicly available cross-linking immunoprecipitation (CLIP)-seq datasets from 51 RBPs.

GREGOR evaluates the enrichment of QTL variants in these genomic annotations by estimating the significance of observed overlap of the eSNP or sSNP relative to the expected overlap using a set of matched control variants. GREGOR creates a list of possible causal SNPs by extending the list of eSNPs or sSNPs (index SNPs) to all SNPs in high linkage disequilibrium (r^2^>0.7). A set of matched control SNPs (SNPs are selected based on matching the index SNP for number of variants in LD, minor allele frequency, and distance to nearest gene/intron) is then created, and enrichments are calculated based on the observed and expected overlap within each annotation.

### eQTL sQTL Overlap

We used the Storey’s *π*_1_, statistic described in Nica et al (Nica et al., 2011), to assess the proportion of true associations among sQTLs that were also detected by the eQTL analysis and eQTLs that were also detected by the sQTL analysis. The overlap was assessed by taking all significant SNP-gene associations from the eQTLs and estimating the proportion of true associations (*π*_1_) on the distribution of corresponding p-values of the overlapping SNP-gene pairs in the sQTL data set and vice versa. This is done by first estimating *π*_0_, the proportion of true null associations based on their distribution. Then *π*_1_=1-*π*_0_ estimates the lower bound of true positive associations.

### Estimation of Variant Effect of eQTL and sQTL

Ensembl’s Variant Effect Predictor (VEP) version 90 (McLaren et al., 2016) was used to annotate the effects of variants of significant QTLs on genes, transcripts, protein sequence, and regulatory regions. VEP annotations are based off of a wide range of reference data including Ensembl database verion 92, GRCH37.p13 genome assembly, Gencode 19 gene annotations, RefSeq 2015-01, PolyPhen 2.2.2, SOFT 5.2.2, dbSNP 150, COSMIC 81, ClinVar 2017-06, and gnomAD r2.0.

### Fetal Cell Type Markers

Cell type enriched genes were obtained from (Polioudakis et al., 2018) single-cell RNA-seq dataset of GW17-18 human fetal cortex. Briefly, Drop-seq was run on single cells isolated from human fetal neocortex according to the online Drop-seq protocol v.3.1 (http://mccarrolllab.com/download/905/) and the methods published in Macosko et al. (Macosko et al., 2015). The raw Drop-seq data was processed using the Drop-seq tools v1.12 pipeline from the McCarroll Laboratory (http://mccarrolllab.com/wp-content/uploads/2016/03/Drop-seqAlignmentCookbookv1.2Jan2016.pdf). Normalization was performed using Seurat v2.0.1 (Butler, Hoffman, Smibert, Papalexi, & Satija, 2018). Raw counts were read depth normalized by dividing by the total number of UMIs per cell, then multiplying by 10,000, adding a value of 1, and log transforming (ln (transcripts-per-10,000 + 1)) using the Seurat function ‘CreateSeuratObject’. To identify cell type enriched genes, differential expression analysis was performed for each cluster individually versus all other cells in the dataset for genes detected in at least 10% of cells in the cluster. Differential expression analysis was performed using a linear model implemented in R as follows: lm(expression ~ number_of_UMI + donor + lab_batch). P-values were then Benjamini-Hochberg corrected (Benjamini & Hochberg, 1995). Genes were considered enriched if they were detected in at least 10% of cells in the cluster, >0.2 log2 fold enriched, and Benjamini-Hochberg corrected p-value < 0.05. Cell type enriched genes were annotated based on the gene harboring an eQTL in either the fetal brain dataset or GTEx adult cortex dataset (Consortium et al., 2017).

### Cross age-tissue comparison

We downloaded GTEx v7 eQTL summary statistics for all 48 tissue types (Consortium et al., 2017). To compare effect sizes consistently between studies, we calculated effect size by running a linear model with scaled log tpm expression values for significant fetal eQTLs and calculated the same way for corresponding SNP-gene pairs in the GTEx data to obtain a beta value from non-standard normalized expression. Significant fetal eQTLs were identified as fetal specific if the corresponding SNP-gene pair was not found in any GTEx tissue or as shared if it was found in at least one GTEx tissue.

Effect size correlations between tissues were calculated by first obtaining all FDR<=0.05 (q-value) nominal fetal eQTLs. Nominal eQTL analysis was run in FastQTL, using the same input for the permutation pass, to obtain all SNP-gene pairs tested. Spearman’s *ρ* correlations were calculated per tissue on the absolute value of the slope from FastQTL output of all FDR<=0.05 fetal eQTLs and corresponding absolute value of slope from SNP-gene pairs in GTEx nominal associations. The absolute value of the slope was used for all correlations to control for strand-flips.

Additionally, we used Storey’s Qvalue software (Storey & Tibshirani, 2003) to assess overlap between fetal brain eQTLs and the eQTLs from the GTEx v7 tissues (Consortium et al., 2017). The proportion of true associations(*π*_1_) was estimated by looking up significant fetal brain eQTLs in each of the GTEx tissues, creating a distribution of corresponding p-values of the overlapping SNP-gene pairs used to calculate π0, the proportion of true null associations based on their distribution. Then *π*_1_=1-*π*_0_ estimates the lower bound of true positive associations. We performed the reciprocal overlap by looking up GTEx significant eQTLs per tissue in the fetal brain dataset (**Figure S6**)

### Partitioned Heritability

Partitioned heritability was measured using LD Score Regression v1.0.0 (Finucane et al., 2015) to identify enrichment of GWAS summary statistics among functional genomic annotations by accounting for LD, specifically eQTL regulatory regions. The full baseline model of 53 functional categories was downloaded from Finucane et al. (https://github.com/bulik/ldsc/wiki/Partitioned-Heritability). Fetal and Adult specific eQTL annotations were created by taking a 500bp window (+-250) around each eSNP and removing any 500bp window that overlapped, resulting in 6163 fetal specific regions and 5690 adult specific regions. An annotation file was then created by marking all HapMap3 (International HapMap, 2003) SNPs that fell within the eQTL annotations. LD scores were calculated for the eQTL annotation SNPs using an LD window of 1cM using LD reference panel 1000 Genomes European Phase 3 (Genomes Project et al., 2015). This LD reference panel was chosen due to the PGC SCZ GWAS (Schizophrenia Working Group of the Psychiatric Genomics, 2014) being comprised of mainly European ancestry samples (Schizophrenia Working Group of the Psychiatric Genomics, 2014). Baseline LD-scores and eQTL LD-scores were simultaneously included in computation of partitioned heritability. Enrichment for each annotation was calculated by the proportion of heritability explained by each annotation divided by the proportion on SNPs in the genome falling in that annotation category. Enrichment p-values were then Bonferroni corrected.

### WGCNA

After gene counts were put through quality control removing genes that were not expressed in 80% of samples with 10 counts or more, expression was conditional quantile normalized, adjusting for gene length and GC content. Sample outliers were removed based on standardized sample network connectivity Z scores < 2. ComBat batch correction was performed (W. E. Johnson et al., 2007). Gestation week, RIN, and the top 4 Picard PCs were regressed from the expression dataset.

Network analysis was performed with robust consensus WGCNA (rWGCNA) (B. Zhang & Horvath, 2005) assigning genes to specific modules based on biweight midcorrelations among genes. Soft threshold power of 11 was chosen to achieve scale-free topology (r2>0.8) (**Figure S8A-B**). Then, 50 signed co-expression networks were generated on 50 independent bootstraps of the samples; each co-expression network uses the same estimated power parameter. The 50 topological overlap matrices were combined edge-wise by taking the median of each edge across all bootstraps. The topological overlap matrices were clustered hierarchically using average linkage hierarchical clustering (using ‘1 – TOM’ as a dis-similarity measure). The topological overlap dendrogram was used to define modules using minimum module size of 100, deep split of 4, merge threshold of 0.2, and negative pamStage.

### GO enrichment

GO definitions were downloaded from Ensembl release 86. GO terms with a small (<35) or large (>100) number of genes were removed. Logistic regression was performed using the model: is.go ~ is.module + gene covariates (GC content and gene length) for an indicator-based enrichment, and p-values were Bonferroni FDR corrected. The top two significant terms are reported.

### Cell Type enrichment

Cell type markers from human and mouse brain were downloaded from (Hawrylycz et al., 2015; Lein et al., 2007; Mancarci et al., 2017; Miller et al., 2014; Tasic et al., 2016; Winden et al., 2009; Y. Zhang et al., 2014; Y. Zhang et al., 2016) as well as the fetal types from (Polioudakis et al., 2018). Logistic regression was performed using the model: is.cell type ~ is.module + gene covariates (GC content and gene length) for an indicator-based enrichment, and p-values were Bonfefrroni FDR corrected. The top two significant cell types are reported.

### Rare variant enrichment

Genes containing disease associated *de novo* variants were downloaded from denovo-db for Autism Spectrum Disorder, Developmental Disorder, Intellectual Disability, and Schizophrnenia (http://denovo-db.gs.washington.edu;(T. N. Turner et al., 2017). Logistic regression was performed using the model: is.disease ~ is.module + gene covariates (GC content and gene length) for an indicator-based enrichment, and p-values were Bonferroni FDR corrected.

### GWAS Module Enrichment

To assess module enrichment of GWAS by gene regulatory regions, we first created a map of gene specific regulatory regions. Every eGene was mapped to its eSNP (1st and 2nd if it had one) and extended to the LD block (r2 >0.8 PLINK) calculated in our sample. GWAS p-values for ASD (Grove et al., 2017) and SCZ (Schizophrenia Working Group of the Psychiatric Genomics, 2014) were assigned to modules based on overlapping positions with eSNP LD block annotations. To assess significance, 1000 permutations were performed for each module, randomly selecting the number of annotations corresponding to the number of genes in each module from all eSNP LD block annotations (**Figure S8E-G**). Significance was calculated by the proportion of permuted observed p-values at the expected p-value of 0.001 that are larger than the actual module’s observed p-value at the expected p-value of 0.001.

### TWAS

We performed SCZ and intercranial volume TWASs using the FUSION package (http://gusevlab.org/projects/fusion/) with gene and splice expression measured in fetal brain tissue. To identify genes with evidence of genetic control, we used GCTA (J. Yang, Lee, Goddard, & Visscher, 2011) software to estimate cis-SNP heritability *h^2g^* (+-1MB window around gene TSS). We identified 3,784 genes and 5,738 splicing-events with significant cis-*h^2g^* (nominal p-value <0.05), which were used to calculate the SNP-based predictive weights per gene/intron.

Using the FUSION package, five-fold cross-validation of five models of expression prediction (best cis-eQTL, best linear unbiased predictor, Bayesian sparse linear mixed model (BSLMM) (X. Zhou, Carbonetto, & Stephens, 2013), Elastic-net regression, LASSO regression) were calculated and evaluated for accuracy. The model with largest cross-validation R^2^ was chosen for downstream association analyses.

TWAS statistics were calculated using fetal weights, as well as published adult expression weights (Gusev et al., 2018)calculated from the CommonMind Consortium eQTL dataset (Fromer et al., 2016), LD SNP correlations from the 1000 Genomes European Phase 3 reference panel (as the GWAS used are from European populations) (Genomes Project et al., 2015), and GWAS summary statistics from PGC2 SCZ GWAS (79,845 individuals) (Schizophrenia Working Group of the Psychiatric Genomics, 2014) and CHARGE ENIGMA Meta-Analysis ICV GWAS (26,577 individuals) (Adams et al., 2016). TWAS association statistics were Bonferroni corrected per GWAS, gene and intron separately.

Overlap of TWAS hits (+-500kb) with GWAS significant loci (LD block reported in paper) was assessed for both fetal TWAS hits and CommonMind Consortium adult brain TWAS hits. Novel regions were identified by genes and introns (+-500k) that did not overlap a GWAS significant loci, and then regions were merged if the gene/intron +-500kb overlapped.

### FOCUS fine mapping

TWAS association statistics at genomic risk regions tend to be correlated as a function of linkage and eQTL overlap between predictive models (Mancuso et al., 2017; Wainberg et al., 2017). To prioritize candidate susceptibility genes in our TWAS we performed statistical fine-mapping using FOCUS (Mancuso et al., 2017). FOCUS models the correlation structure induced by LD and overlapping eQTL weights across predictive models and computes posterior probabilities for a gene/intron to explain all observed TWAS association signal at a region. We next computed 90% credible gene-sets by taking genes with largest posterior probability until 90% density was explained.

## Supplemental Information

**Figure S1.**
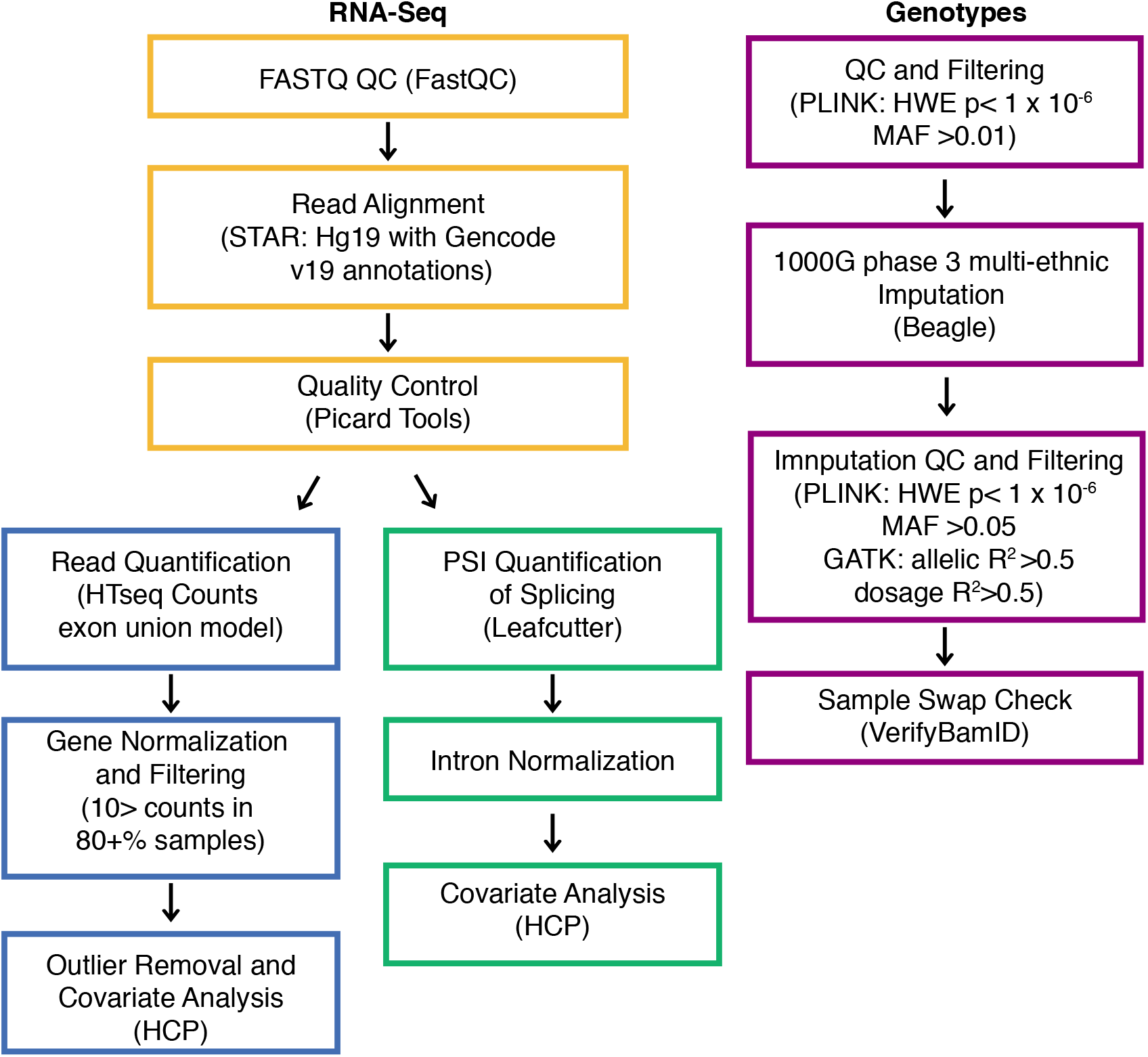
Related to Figure 1 and Experimental Procedures: Overview of methods and QC pipeline for processing RNA sequencing FASTQ files to gene and intron quantifications and genotype imputation, QC, and filtering. Output data from this pipeline was used as inputs for further eQTL and sQTL analysis.

**Figure S2.**
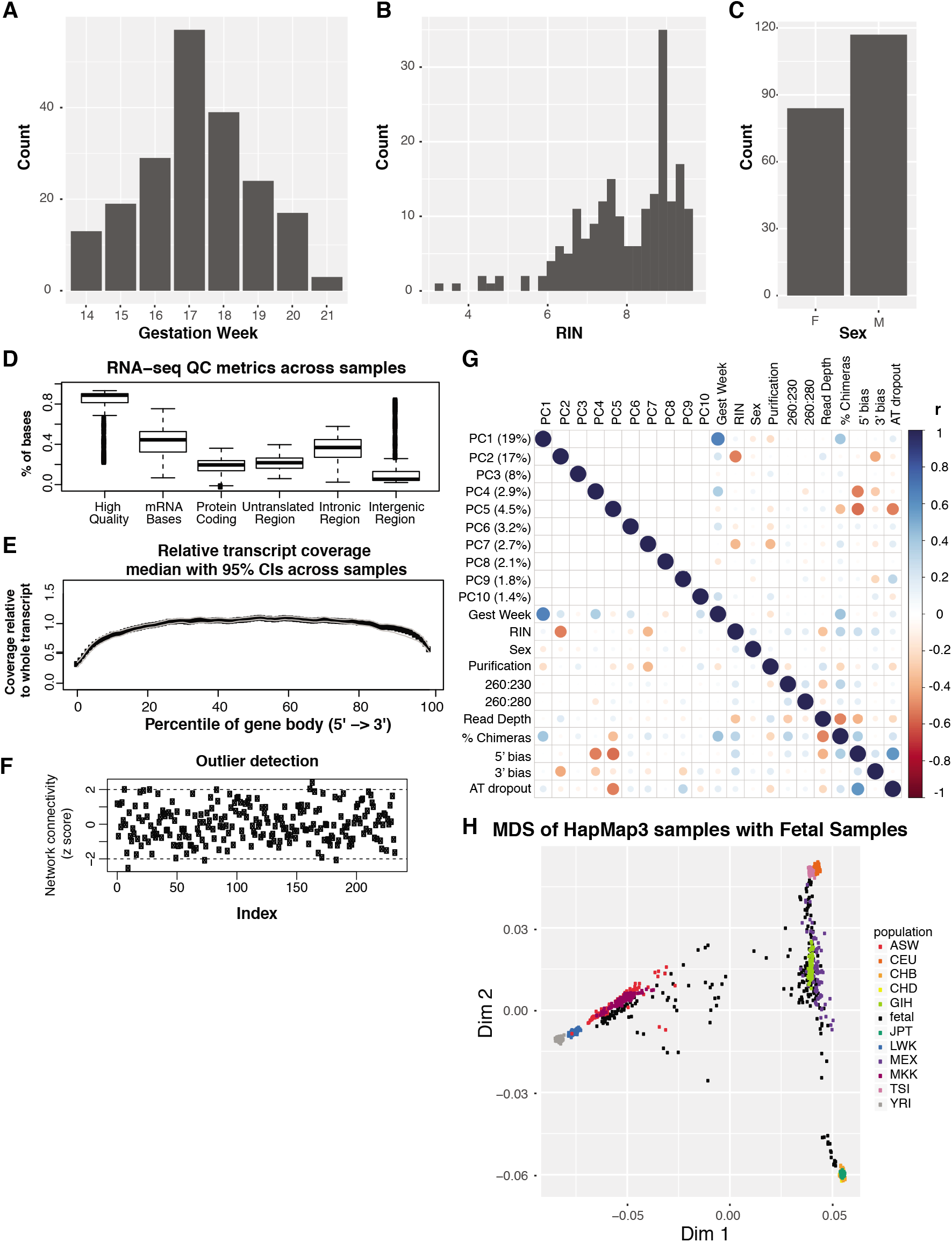
Related to Experimental Procedures: Sample demographics and dataset quality control. **(A)** Distribution of age (gestation week) of our samples, showing our samples span midgestation. **(B)** Distribution of RNA integrity number (RIN) among our samples. **(C)** Distribution of sex among our samples. **(D)** RNA sequencing metrics from Picard tools showing the majority of our reads are of high quality and correspond to mRNA. **(E)** Relative transcript coverage across the body of a gene, showing good coverage along the entire gene body, representative of using a ribo-zero library preparation. **(F)** Sample outlier detection determined from WGCNA’s network connectivity Z-score. Samples with a Z-score of greater than 2 or less than -2 are removed. **(G)** Correlation of the top 10 expression principle components (PCs) with gestation week, RIN, sex, Picard tool metrics: read depth, % chimeras, 5’ bias, 3’ bias, AT dropout, and technical measurements: purification method and purity assessments 260:230 and 260:280. Gestation Week and RIN show high correlations with the top 2 PCs. **(H)** MDS plot of genotypes from our fetal brain samples merged with the HapMap3 samples from 11 populations, shows the diversity across the fetal samples.

**Figure S3.**
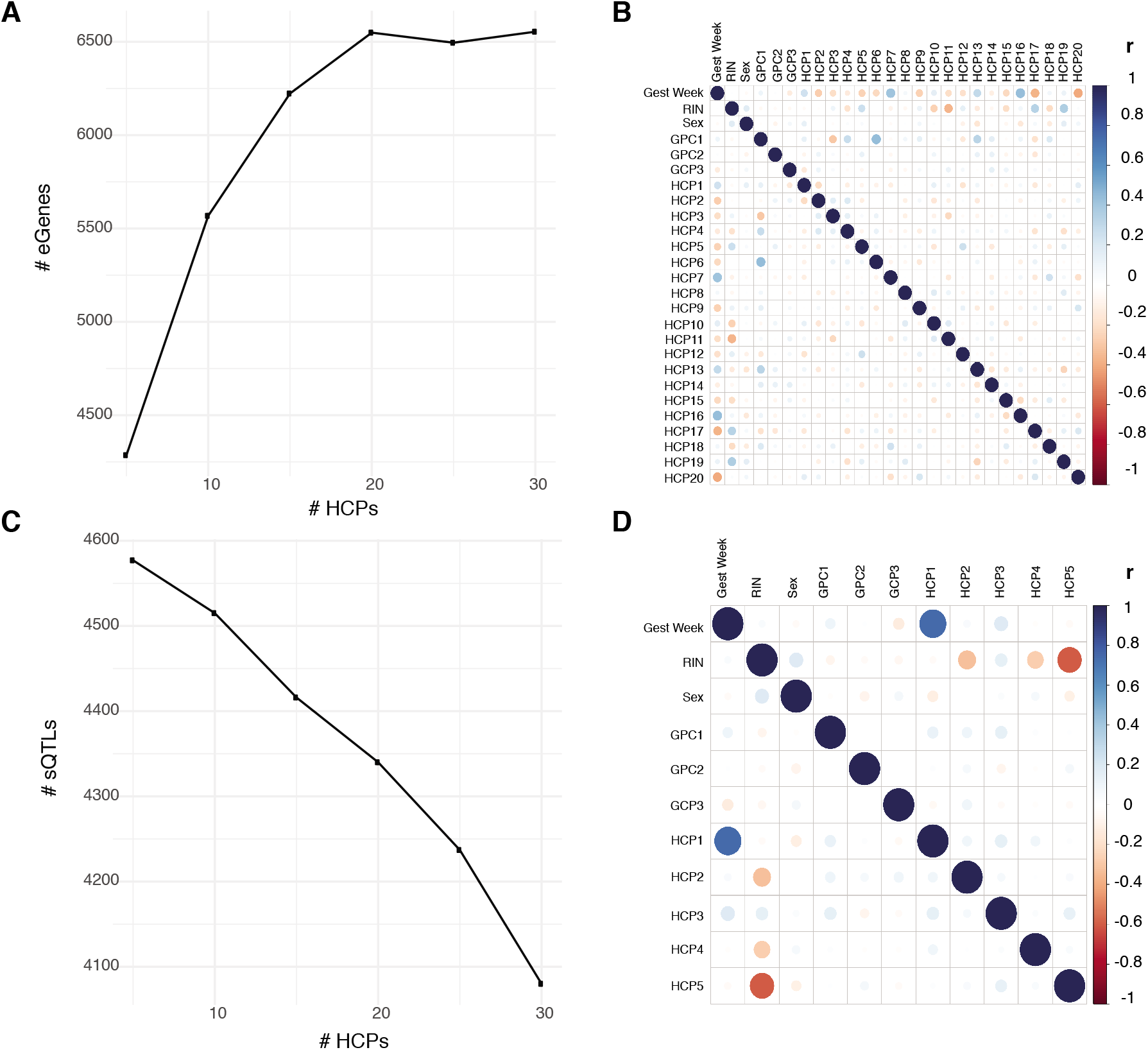
Related to Experimental Procedures: Identification of optimal number of HCP selection and correlations of all covariates in final cis-eQTL analysis. **(A)** The number of hidden covariates (HCP) was chosen to maximize cis-eGene discovery. Cis-eQTL analysis was performed correcting for gestation week, RIN, sex, the top 3 genotype PCs and then increments of 5 HCPs using FastQTLs permutation scheme. The number of HCPs included in the eQTL analysis is shown on the X axis with the number of significant eGenes is shown on the Y axis. The number of eGenes was determined by FDR q-value <=0.05. Based on these results, 20 HCPs were selected for the final eQTL analysis. **(B)** Pearson correlation (r) between all covariates used in the final eQTL analysis. **(C)** The number of HCPs chosen to maximize sQTL discovery. sQTL analysis was performed correcting for gestation week, RIN, sex, the top 3 genotype PCs and then increments of 5 HCPs using FastQTLs permutation scheme. The number of HCPs included in the sQTL analysis is shown on the X axis with the number of significant sQTLs is shown on the Y axis. The number of sQTLs (top SNP per intron) was determined by FDR q-value <=0.05. Based on these results, 5 HCPs were selected for the final sQTL analysis. **(D)** Pearson correlation (r) between all covariates used in the final sQTL analysis.

**Figure S4.**
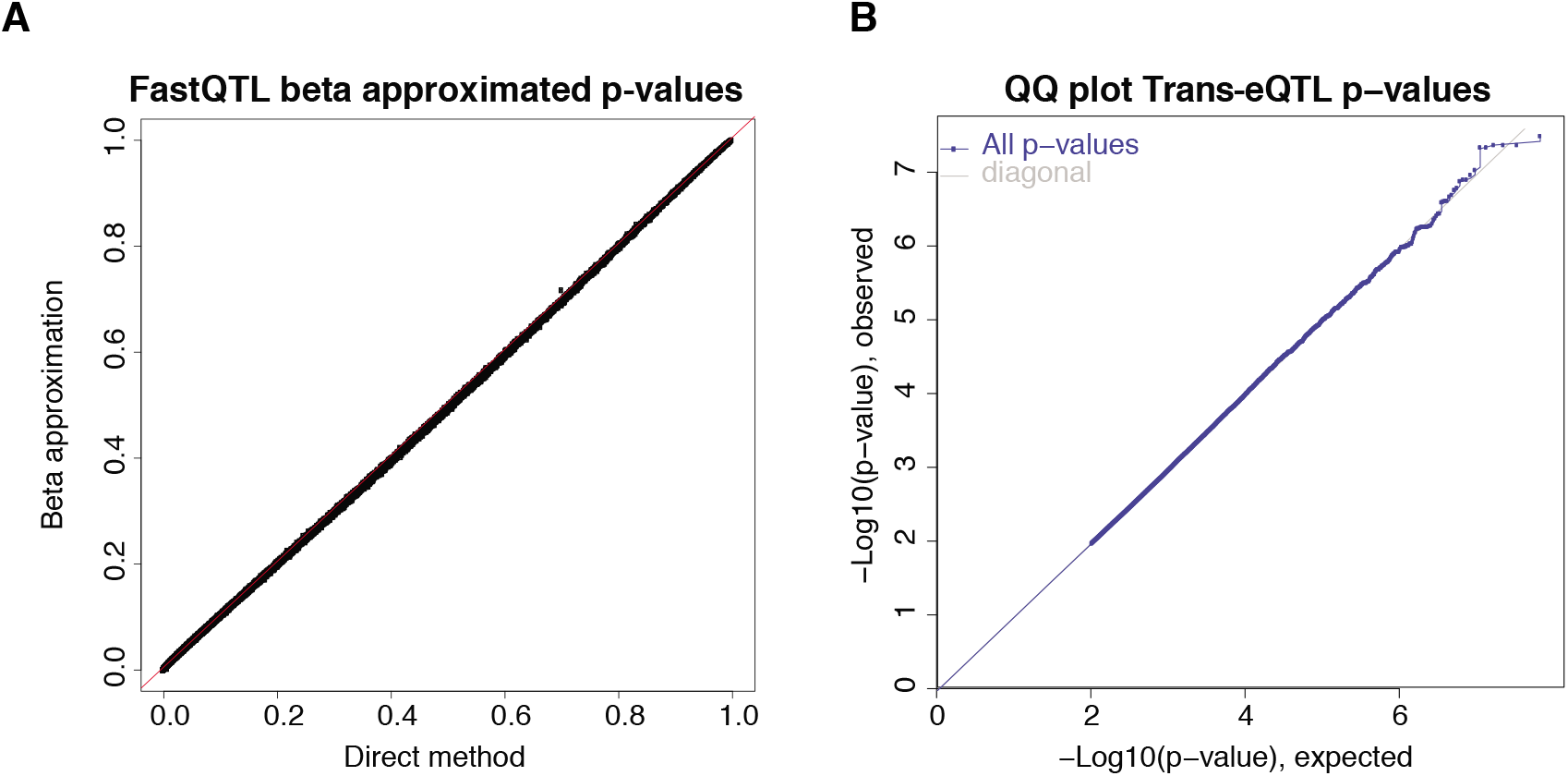
Related to Experimental Procedures: FastQTL QC. **(A)** FastQTL implements a beta approximation for permutation p-values. Empirical p-values are on the X axis with beta approximated p-values are on the Y axis. This shows the beta approximated p-values are well calibrated given the empirical p-values. **(B)** QQ plot for trans-eQTL analysis that was run for 10 randomly chosen genes, expected p-values on the X axis and observed p-values on the Y axis, indicating there is no systematic inflation.

**Figure S5.**
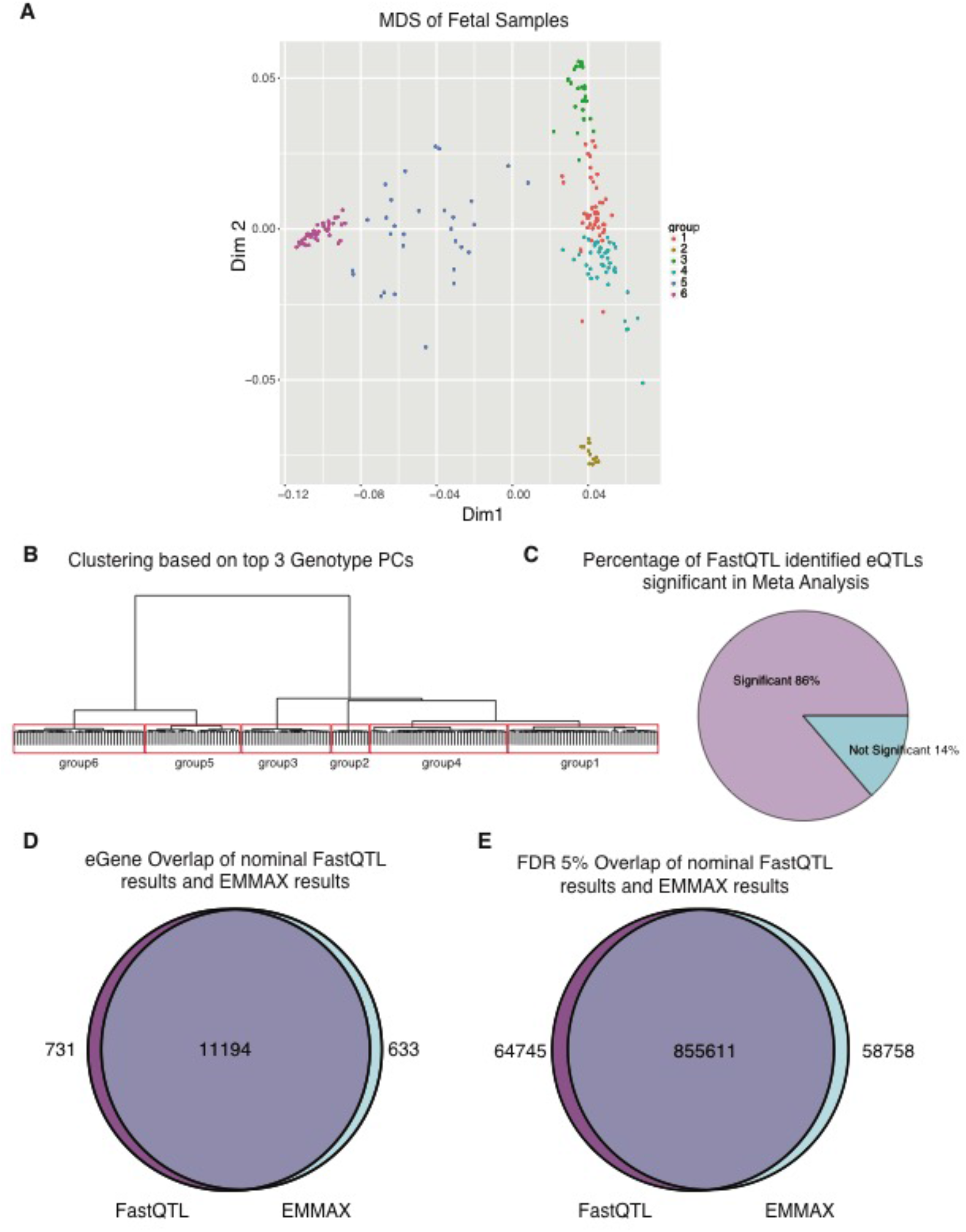
Related to Experimental Procedures: Ancestry correction. **(A)** MDS plot of genotypes from just fetal brain samples colored by group assignments from (B). **(B)** Sample clustering based on hierarchical clustering of the top 3 genotype PCs groups fetal brain samples into 6 subpopulation groups. **(C)** Meta-Analysis of eQTLs (top SNP per gene) performed across the 6 subpopulation groups finds 86% of eQTLs significant. **(D)** 94% of eGenes in FastQTLs nominal pass overlap significant eGenes from EMMAX. **(E)** 93% of significant (FDR <=0.05) SNP-gene pairs from the nominal FastQTL pass overlap with significant SNP-gene pairs from the EMMAX eQTL analysis.

**Figure S6.**
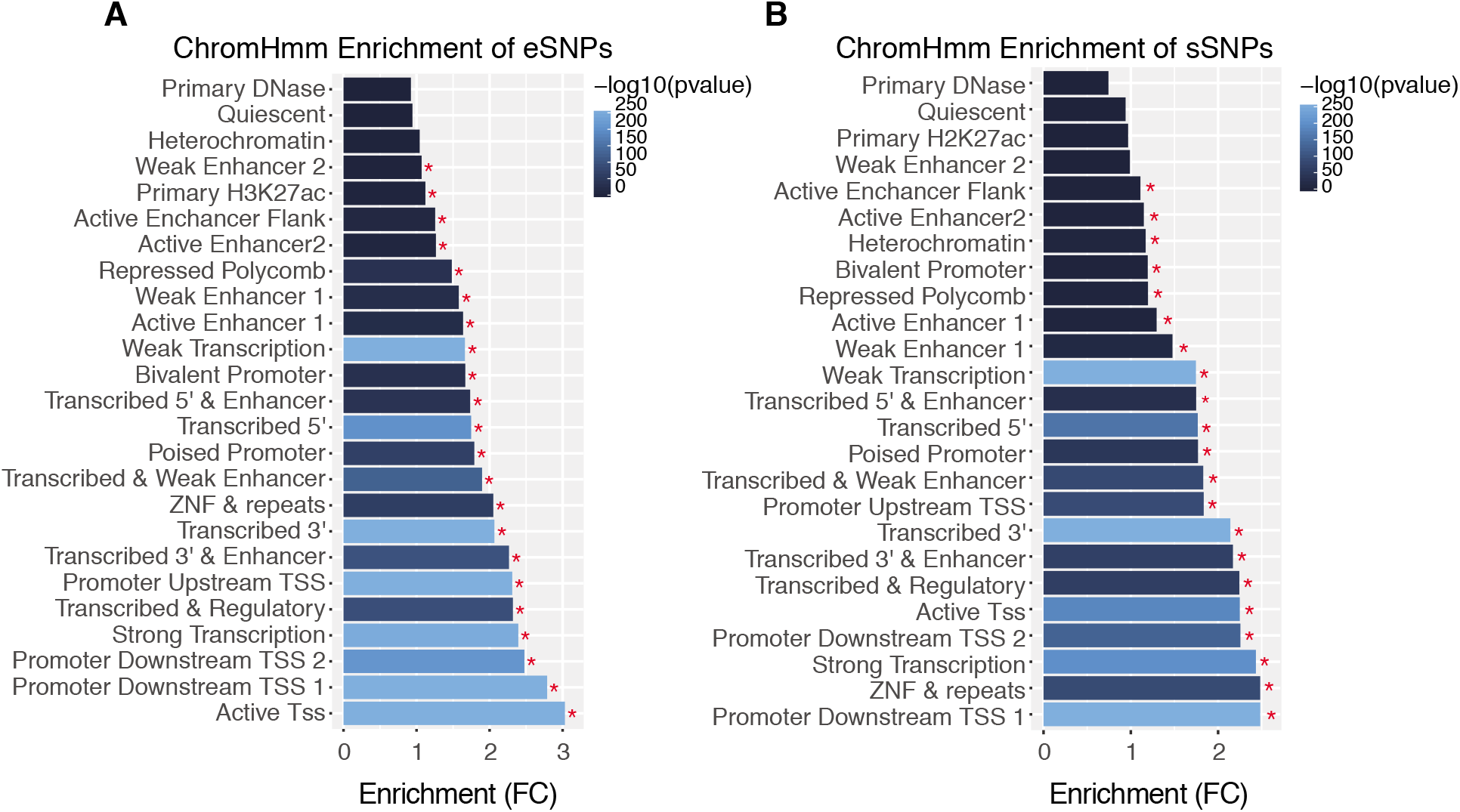
Related to Figure 2F and Figure 3C: Full ChromHmm Model **(A-B)** Enrichments of eQTLs and sQTLs in the full ChromHmm model of 25 states. Fold enrichment of e/sSNPs by their distribution within specific fetal brain chromatin states, which are depicted on the Y axis (ChromHMM annotations; Methods)

**Figure S7.**
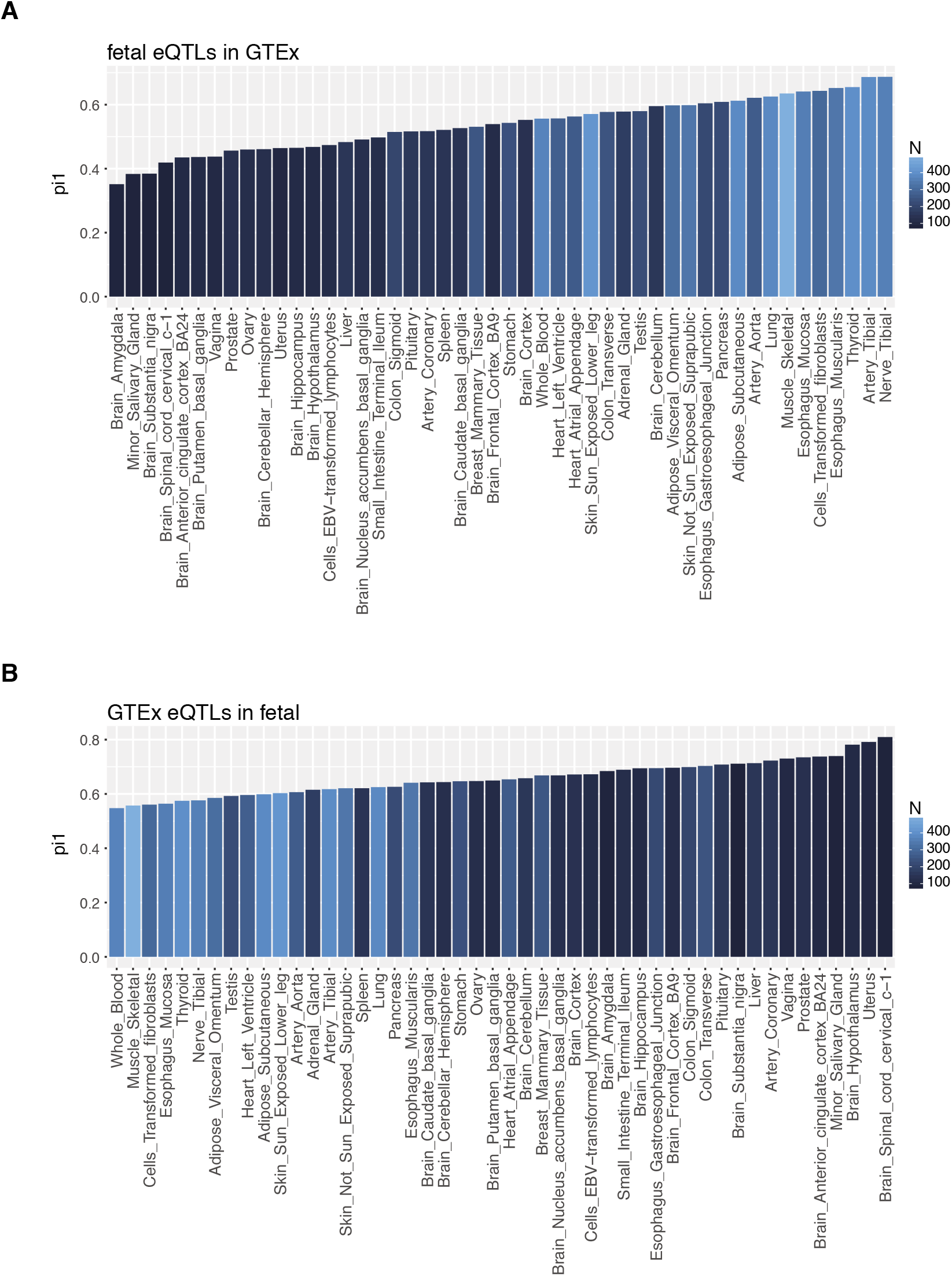
Related to Figure 4: P-Value based tissue overlap are a function of sample size. **(A)** Pi1 values (proportion of true positive p-values) for significant fetal brain eQTLs in GTEx tissues, colored by sample size (N) of GTEx tissue dataset. **(B)** Pi1 values (proportion of true positive p-values) for significant GTEx eQTLs per tissue in the fetal brain eQTLs, colored by sample size (N) of GTEx tissue dataset.

**Figure S8.**
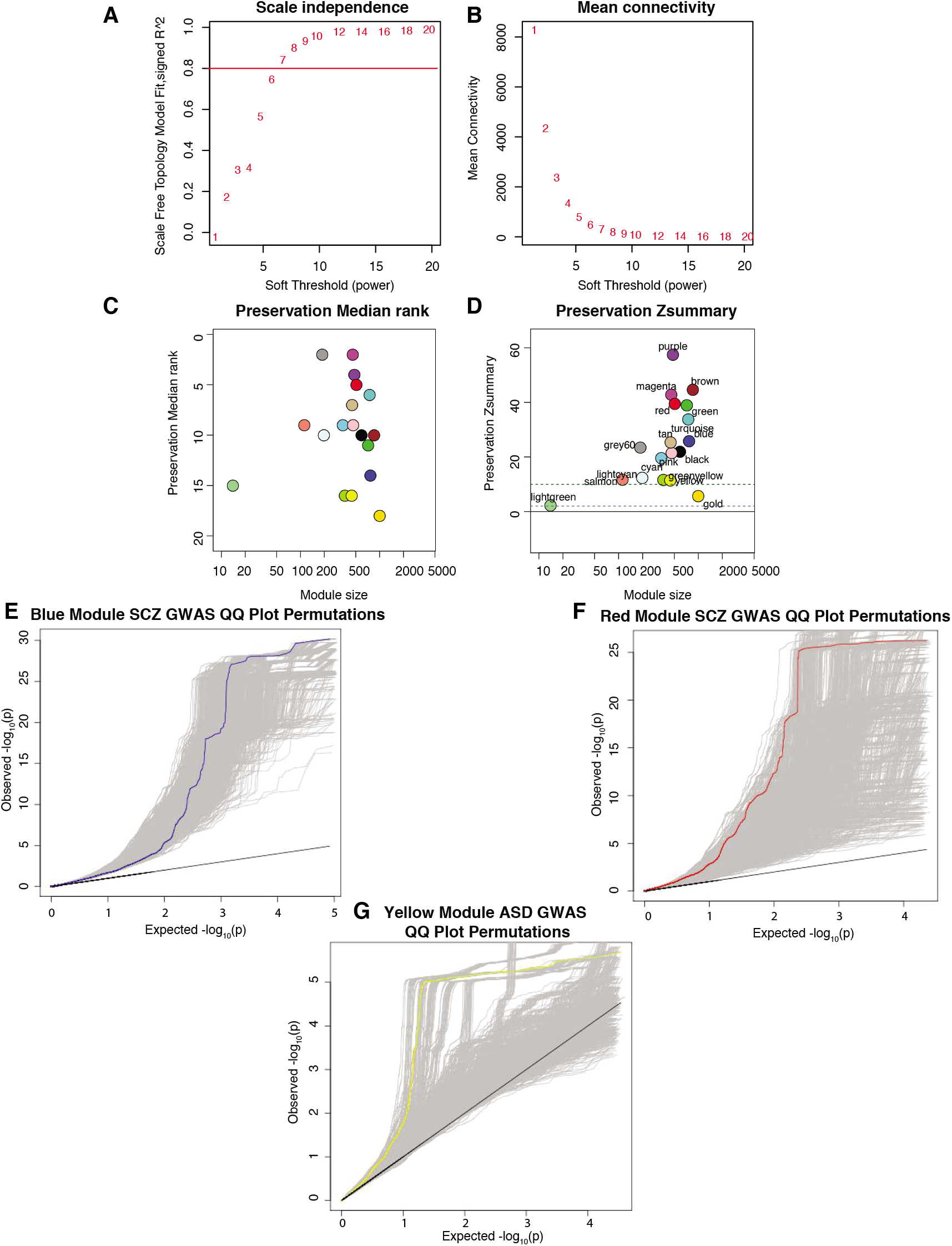
Related to Figure 5: WGCNA **(A-B)** Soft threshold power of 11 was chosen based on scale free topology plateauing at a soft threshold power of 11, as well as mean connectivity. **(C-D)** Preservation median rand and Z summary of fetal brain modules in the (Parikshak et al., 2013) network. **(E-G)** Per module QQ plot of GWAS SNP p-values (SCZ GWAS for blue and red module, ASD GWAS for yellow module) in regulatory regions defined by eQTLs, with 1000 permutations in grey. Expected p-values shown on the X axis and observed p-values shown on the Y axis.

## References

Adams, H. H., Hibar, D. P., Chouraki, V., Stein, J. L., Nyquist, P. A., Renteria, M. E., … Thompson, P. M. (2016). Novel genetic loci underlying human intracranial volume identified through genome-wide association. Nat Neurosci, 19(12), 1569–1582. doi:10.1038/nn.4398

Almaguer-Mederos, L. E., Mesa, J. M. L., Gonzalez-Zaldivar, Y., Almaguer-Gotay, D., Cuello-Almarales, D., Aguilera-Rodriguez, R., … Velazquez-Perez, L. (2018). Factors associated with ATXN2 CAG/CAA repeat intergenerational instability in Spinocerebellar ataxia type 2. Clin Genet. doi:10.1111/cge.13380

Anders, S., Pyl, P. T., & Huber, W. (2015). HTSeq--a Python framework to work with high-throughput sequencing data. Bioinformatics, 31(2), 166–169. doi:10.1093/bioinformatics/btu638

Arbiza, L., Gronau, I., Aksoy, B. A., Hubisz, M. J., Gulko, B., Keinan, A., & Siepel, A. (2013). Genome-wide inference of natural selection on human transcription factor binding sites. Nat Genet, 45(7), 723–729. doi:10.1038/ng.2658

Autism Spectrum Disorders Working Group of The Psychiatric Genomics, C. (2017). Meta-analysis of GWAS of over 16,000 individuals with autism spectrum disorder highlights a novel locus at 10q24.32 and a significant overlap with schizophrenia. Mol Autism, 8, 21. doi:10.1186/s13229-017-0137-9

Bae, B. I., Jayaraman, D., & Walsh, C. A. (2015). Genetic changes shaping the human brain. Dev Cell, 32(4), 423–434. doi:10.1016/j.devcel.2015.01.035

Battle, A., Khan, Z., Wang, S. H., Mitrano, A., Ford, M. J., Pritchard, J. K., & Gilad, Y. (2015). Genomic variation. Impact of regulatory variation from RNA to protein. Science, 347(6222), 664–667. doi:10.1126/science.1260793

Benjamini, Y., & Hochberg, Y. (1995). Controlling the false discovery rate: a practical and powerful approach to multiple testing. J Roy Stat Soc B Met, 57, 289–300.

Bernier, R., Hudac, C. M., Chen, Q., Zeng, C., Wallace, A. S., Gerdts, J., … Simons, V. I. P. c. (2017). Developmental trajectories for young children with 16p11.2 copy number variation. Am J Med Genet B Neuropsychiatr Genet, 174(4), 367–380. doi:10.1002/ajmg.b.32525

Bestman, J. E., Huang, L. C., Lee-Osbourne, J., Cheung, P., & Cline, H. T. (2015). An in vivo screen to identify candidate neurogenic genes in the developing Xenopus visual system. Dev Biol, 408(2), 269–291. doi:10.1016/j.ydbio.2015.03.010

BrainSpan. (2013). BrainSpan: Atlas of the Developing Human Brain. Retrieved from http://www.brainspan.org

Browning, B. L., & Browning, S. R. (2016). Genotype Imputation with Millions of Reference Samples. Am J Hum Genet, 98(1), 116–126. doi:10.1016/j.ajhg.2015.11.020

Butler, A., Hoffman, P., Smibert, P., Papalexi, E., & Satija, R. (2018). Integrating single-cell transcriptomic data across different conditions, technologies, and species. Nat Biotechnol, 36(5), 411–420. doi:10.1038/nbt.4096

Carvill, G. L., Heavin, S. B., Yendle, S. C., McMahon, J. M., O’Roak, B. J., Cook, J., … Mefford, H. C. (2013). Targeted resequencing in epileptic encephalopathies identifies de novo mutations in CHD2 and SYNGAP1. Nat Genet, 45(7), 825–830. doi:10.1038/ng.2646

Chang, C. C., Chow, C. C., Tellier, L. C., Vattikuti, S., Purcell, S. M., & Lee, J. J. (2015). Second-generation PLINK: rising to the challenge of larger and richer datasets. Gigascience, 4, 7. doi:10.1186/s13742-015-0047-8

Choi, J., Ko, J., Racz, B., Burette, A., Lee, J. R., Kim, S., … Kim, E. (2005). Regulation of dendritic spine morphogenesis by insulin receptor substrate 53, a downstream effector of Rac1 and Cdc42 small GTPases. J Neurosci, 25(4), 869–879. doi:10.1523/JNEUROSCI.3212-04.2005

Cockerill, P. N. (2011). Structure and function of active chromatin and DNase I hypersensitive sites. FEBS J, 278(13), 2182–2210. doi:10.1111/j.1742-4658.2011.08128.x

Colantuoni, C., Lipska, B. K., Ye, T., Hyde, T. M., Tao, R., Leek, J. T., … Kleinman, J. E. (2011). Temporal dynamics and genetic control of transcription in the human prefrontal cortex. Nature, 478(7370), 519–523. doi:10.1038/nature10524

Consortium, G. T. (2015). Human genomics. The Genotype-Tissue Expression (GTEx) pilot analysis: multitissue gene regulation in humans. Science, 348(6235), 648–660. doi:10.1126/science.1262110

Consortium, G. T., Laboratory, D. A., Coordinating Center-Analysis Working, G., Statistical Methods groups-Analysis Working, G., Enhancing, G. g., Fund, N. I. H. C., … Montgomery, S. B. (2017). Genetic effects on gene expression across human tissues. Nature, 550(7675), 204–213. doi:10.1038/nature24277

Darvish, H., Azcona, L. J., Tafakhori, A., Ahmadi, M., Ahmadifard, A., & Paisan-Ruiz, C. (2017). Whole genome sequencing identifies a novel homozygous exon deletion in the NT5C2 gene in a family with intellectual disability and spastic paraplegia. NPJ Genom Med, 2. doi:10.1038/s41525-017-0022-7

de la Torre-Ubieta, L., Stein, J. L., Won, H., Opland, C. K., Liang, D., Lu, D., & Geschwind, D. H. (2018). The Dynamic Landscape of Open Chromatin during Human Cortical Neurogenesis. Cell, 172(1–2), 289–304 e218. doi:10.1016/j.cell.2017.12.014

Dobin, A., Davis, C. A., Schlesinger, F., Drenkow, J., Zaleski, C., Jha, S., … Gingeras, T. R. (2013). STAR: ultrafast universal RNA-seq aligner. Bioinformatics, 29(1), 15–21. doi:10.1093/bioinformatics/bts635

Durak, O., Gao, F., Kaeser-Woo, Y. J., Rueda, R., Martorell, A. J., Nott, A., … Tsai, L. H. (2016). Chd8 mediates cortical neurogenesis via transcriptional regulation of cell cycle and Wnt signaling. Nat Neurosci, 19(11), 1477–1488. doi:10.1038/nn.4400

Durinck, S., Spellman, P. T., Birney, E., & Huber, W. (2009). Mapping identifiers for the integration of genomic datasets with the R/Bioconductor package biomaRt. Nat Protoc, 4(8), 1184–1191. doi:10.1038/nprot.2009.97

Eising, E., Carrion-Castillo, A., Vino, A., Strand, E. A., Jakielski, K. J., Scerri, T. S., … Fisher, S. E. (2018). A set of regulatory genes co-expressed in embryonic human brain is implicated in disrupted speech development. Mol Psychiatry. doi:10.1038/s41380-018-0020-x

Ernst, J., & Kellis, M. (2015). Large-scale imputation of epigenomic datasets for systematic annotation of diverse human tissues. Nat Biotechnol, 33(4), 364–376. doi:10.1038/nbt.3157

Ernst, J., Kheradpour, P., Mikkelsen, T. S., Shoresh, N., Ward, L. D., Epstein, C. B., … Bernstein, B. E. (2011). Mapping and analysis of chromatin state dynamics in nine human cell types. Nature, 473(7345), 43–49. doi:10.1038/nature09906

Escamilla, C. O., Filonova, I., Walker, A. K., Xuan, Z. X., Holehonnur, R., Espinosa, F., … Powell, C. M. (2017). Kctd13 deletion reduces synaptic transmission via increased RhoA. Nature, 551(7679), 227–231. doi:10.1038/nature24470

Fan, J., Zhou, X., Wang, Y., Kuang, C., Sun, Y., Liu, X., … Xu, Y. (2017). Differential requirement for N-ethylmaleimide-sensitive factor in endosomal trafficking of transferrin receptor from anterograde trafficking of vesicular stomatitis virus glycoprotein G. FEBS Lett, 591(2), 273–281. doi:10.1002/1873-3468.12532

Finucane, H. K., Bulik-Sullivan, B., Gusev, A., Trynka, G., Reshef, Y., Loh, P. R., … Price, A. L. (2015). Partitioning heritability by functional annotation using genome-wide association summary statistics. Nat Genet, 47(11), 1228–1235. doi:10.1038/ng.3404

Flutre, T., Wen, X., Pritchard, J., & Stephens, M. (2013). A statistical framework for joint eQTL analysis in multiple tissues. PLoS Genet, 9(5), e1003486. doi:10.1371/journal.pgen.1003486

Fromer, M., Roussos, P., Sieberts, S. K., Johnson, J. S., Kavanagh, D. H., Perumal, T. M., … Sklar, P. (2016). Gene expression elucidates functional impact of polygenic risk for schizophrenia. Nat Neurosci, 19(11), 1442–1453. doi:10.1038/nn.4399

Gandal, M. J., Leppa, V., Won, H., Parikshak, N. N., & Geschwind, D. H. (2016). The road to precision psychiatry: translating genetics into disease mechanisms. Nat Neurosci, 19(11), 1397–1407. doi:10.1038/nn.4409

Genomes Project, C., Auton, A., Brooks, L. D., Durbin, R. M., Garrison, E. P., Kang, H. M., … Abecasis, G. R. (2015). A global reference for human genetic variation. Nature, 526(7571), 68–74. doi:10.1038/nature15393

Geschwind, D. H., & Flint, J. (2015). Genetics and genomics of psychiatric disease. Science, 349(6255), 1489–1494. doi:10.1126/science.aaa8954

Geschwind, D. H., & Rakic, P. (2013). Cortical evolution: judge the brain by its cover. Neuron, 80(3), 633–647. doi:10.1016/j.neuron.2013.10.045

Gilman, S. R., Chang, J., Xu, B., Bawa, T. S., Gogos, J. A., Karayiorgou, M., & Vitkup, D. (2012). Diverse types of genetic variation converge on functional gene networks involved in schizophrenia. Nat Neurosci, 15(12), 1723–1728. doi:10.1038/nn.3261

Golzio, C., Willer, J., Talkowski, M. E., Oh, E. C., Taniguchi, Y., Jacquemont, S., … Katsanis, N. (2012). KCTD13 is a major driver of mirrored neuroanatomical phenotypes of the 16p11.2 copy number variant. Nature, 485(7398), 363–367. doi:10.1038/nature11091

Grammatikakis, I., Zhang, P., Panda, A. C., Kim, J., Maudsley, S., Abdelmohsen, K., … Gorospe, M. (2016). Alternative Splicing of Neuronal Differentiation Factor TRF2 Regulated by HNRNPH1/H2. Cell Rep, 15(5), 926–934. doi:10.1016/j.celrep.2016.03.080

Gratten, J., Wray, N. R., Keller, M. C., & Visscher, P. M. (2014). Large-scale genomics unveils the genetic architecture of psychiatric disorders. Nat Neurosci, 17(6), 782–790. doi:10.1038/nn.3708

Grove, J., Ripke, S., Als, T. D., Mattheisen, M., Walters, R., Won, H., … Børglum, A. D. (2017). Common risk variants identified in autism spectrum disorder. bioRxiv. doi:10.1101/224774

Gulsuner, S., Walsh, T., Watts, A. C., Lee, M. K., Thornton, A. M., Casadei, S., … McClellan, J. M. (2013). Spatial and temporal mapping of de novo mutations in schizophrenia to a fetal prefrontal cortical network. Cell, 154(3), 518–529. doi:10.1016/j.cell.2013.06.049

Gusev, A., Ko, A., Shi, H., Bhatia, G., Chung, W., Penninx, B. W., … Pasaniuc, B. (2016). Integrative approaches for large-scale transcriptome-wide association studies. Nat Genet, 48(3), 245–252. doi:10.1038/ng.3506

Gusev, A., Mancuso, N., Won, H., Kousi, M., Finucane, H. K., Reshef, Y., … Price, A. L. (2018). Transcriptome-wide association study of schizophrenia and chromatin activity yields mechanistic disease insights. Nat Genet, 50(4), 538–548. doi:10.1038/s41588-018-0092-1

Hahne, F., & Ivanek, R. (2016). Visualizing Genomic Data Using Gviz and Bioconductor. Methods Mol Biol, 1418, 335–351. doi:10.1007/978-1-4939-3578-9_16

Hannon, E., Spiers, H., Viana, J., Pidsley, R., Burrage, J., Murphy, T. M., … Mill, J. (2016). Methylation QTLs in the developing brain and their enrichment in schizophrenia risk loci. Nat Neurosci, 19(1), 48–54. doi:10.1038/nn.4182

Hansen, K. D., Irizarry, R. A., & Wu, Z. (2012). Removing technical variability in RNA-seq data using conditional quantile normalization. Biostatistics, 13(2), 204–216. doi:10.1093/biostatistics/kxr054

Hanson, E., Bernier, R., Porche, K., Jackson, F. I., Goin-Kochel, R. P., Snyder, L. G., … Simons Variation in Individuals Project, C. (2015). The cognitive and behavioral phenotype of the 16p11.2 deletion in a clinically ascertained population. Biol Psychiatry, 77(9), 785–793. doi:10.1016/j.biopsych.2014.04.021

Harrow, J., Frankish, A., Gonzalez, J. M., Tapanari, E., Diekhans, M., Kokocinski, F., … Hubbard, T. J. (2012). GENCODE: the reference human genome annotation for The ENCODE Project. Genome Res, 22(9), 1760–1774. doi:10.1101/gr.135350.111

Hawrylycz, M., Miller, J. A., Menon, V., Feng, D., Dolbeare, T., Guillozet-Bongaarts, A. L., … Lein, E. (2015). Canonical genetic signatures of the adult human brain. Nat Neurosci, 18(12), 1832–1844. doi:10.1038/nn.4171

Ingason, A., Giegling, I., Hartmann, A. M., Genius, J., Konte, B., Friedl, M., … Rujescu, D. (2015). Expression analysis in a rat psychosis model identifies novel candidate genes validated in a large case-control sample of schizophrenia. Transl Psychiatry, 5, e656. doi:10.1038/tp.2015.151

International HapMap, C. (2003). The International HapMap Project. Nature, 426(6968), 789–796. doi:10.1038/nature02168

Iossifov, I., Ronemus, M., Levy, D., Wang, Z., Hakker, I., Rosenbaum, J., … Wigler, M. (2012). De novo gene disruptions in children on the autistic spectrum. Neuron, 74(2), 285–299. doi:10.1016/j.neuron.2012.04.009

Jaffe, A. E., Gao, Y., Deep-Soboslay, A., Tao, R., Hyde, T. M., Weinberger, D. R., & Kleinman, J. E. (2016). Mapping DNA methylation across development, genotype and schizophrenia in the human frontal cortex. Nat Neurosci, 19(1), 40–47. doi:10.1038/nn.4181

Jaffe, A. E., Straub, R. E., Shin, J. H., Tao, R., Gao, Y., Collado-Torres, L., … Weinberger, D. R. (2018). Developmental and genetic regulation of the human cortex transcriptome illuminate schizophrenia pathogenesis. Nat Neurosci, 21(8), 1117–1125. doi:10.1038/s41593-018-0197-y

Johnson, M. B., Kawasawa, Y. I., Mason, C. E., Krsnik, Z., Coppola, G., Bogdanovic, D., … Sestan, N. (2009). Functional and evolutionary insights into human brain development through global transcriptome analysis. Neuron, 62(4), 494–509. doi:10.1016/j.neuron.2009.03.027

Johnson, W. E., Li, C., & Rabinovic, A. (2007). Adjusting batch effects in microarray expression data using empirical Bayes methods. Biostatistics, 8(1), 118–127. doi:10.1093/biostatistics/kxj037

Jun, G., Flickinger, M., Hetrick, K. N., Romm, J. M., Doheny, K. F., Abecasis, G. R., … Kang, H. M. (2012). Detecting and estimating contamination of human DNA samples in sequencing and array-based genotype data. Am J Hum Genet, 91(5), 839–848. doi:10.1016/j.ajhg.2012.09.004

Kang, H. J., Kawasawa, Y. I., Cheng, F., Zhu, Y., Xu, X., Li, M., … Sestan, N. (2011). Spatio-temporal transcriptome of the human brain. Nature, 478(7370), 483–489. doi:10.1038/nature10523

Kang, H. M., Zaitlen, N. A., Wade, C. M., Kirby, A., Heckerman, D., Daly, M. J., & Eskin, E. (2008). Efficient control of population structure in model organism association mapping. Genetics, 178(3), 1709–1723. doi:10.1534/genetics.107.080101

Kim, Y., Xia, K., Tao, R., Giusti-Rodriguez, P., Vladimirov, V., van den Oord, E., & Sullivan, P. F. (2014). A meta-analysis of gene expression quantitative trait loci in brain. Transl Psychiatry, 4, e459. doi:10.1038/tp.2014.96

Kim, Y. E., Oh, K. W., Noh, M. Y., Park, J., Kim, H. J., Park, J. E., … Kim, S. H. (2018). Analysis of ATXN2 trinucleotide repeats in Korean patients with amyotrophic lateral sclerosis. Neurobiol Aging, 67, 201 e205–201 e208. doi:10.1016/j.neurobiolaging.2018.03.019

Kostovic, I., & Jovanov-Milosevic, N. (2006). The development of cerebral connections during the first 20-45 weeks’ gestation. Semin Fetal Neonatal Med, 11(6), 415–422. doi:10.1016/j.siny.2006.07.001

Lai, C. S., Fisher, S. E., Hurst, J. A., Vargha-Khadem, F., & Monaco, A. P. (2001). A forkhead-domain gene is mutated in a severe speech and language disorder. Nature, 413(6855), 519–523. doi:10.1038/35097076

Langfelder, P., & Horvath, S. (2008). WGCNA: an R package for weighted correlation network analysis. BMC Bioinformatics, 9, 559. doi:10.1186/1471-2105-9-559

Langfelder, P., & Horvath, S. (2012). Fast R Functions for Robust Correlations and Hierarchical Clustering. J Stat Softw, 46(11).

Lappalainen, T., Sammeth, M., Friedlander, M. R., t Hoen, P. A., Monlong, J., Rivas, M. A., … Dermitzakis, E. T. (2013). Transcriptome and genome sequencing uncovers functional variation in humans. Nature, 501(7468), 506–511. doi:10.1038/nature12531

Latourelle, J. C., Dumitriu, A., Hadzi, T. C., Beach, T. G., & Myers, R. H. (2012). Evaluation of Parkinson disease risk variants as expression-QTLs. PLoS One, 7(10), e46199. doi:10.1371/journal.pone.0046199

Lawrence, M., Huber, W., Pages, H., Aboyoun, P., Carlson, M., Gentleman, R., … Carey, V. J. (2013). Software for computing and annotating genomic ranges. PLoS Comput Biol, 9(8), e1003118. doi:10.1371/journal.pcbi.1003118

Leek, J. T., & Storey, J. D. (2007). Capturing heterogeneity in gene expression studies by surrogate variable analysis. PLoS Genet, 3(9), 1724–1735. doi:10.1371/journal.pgen.0030161

Lein, E. S., Hawrylycz, M. J., Ao, N., Ayres, M., Bensinger, A., Bernard, A., … Jones, A. R. (2007). Genome-wide atlas of gene expression in the adult mouse brain. Nature, 445(7124), 168–176. doi:10.1038/nature05453

Lek, M., Karczewski, K. J., Minikel, E. V., Samocha, K. E., Banks, E., Fennell, T., … Exome Aggregation, C. (2016). Analysis of protein-coding genetic variation in 60,706 humans. Nature, 536(7616), 285–291. doi:10.1038/nature19057

Li, H., Handsaker, B., Wysoker, A., Fennell, T., Ruan, J., Homer, N., … Genome Project Data Processing, S. (2009). The Sequence Alignment/Map format and SAMtools. Bioinformatics, 25(16), 2078–2079. doi:10.1093/bioinformatics/btp352

Li, Y. I., Knowles, D. A., Humphrey, J., Barbeira, A. N., Dickinson, S. P., Im, H. K., & Pritchard, J. K. (2018). Annotation-free quantification of RNA splicing using LeafCutter. Nat Genet, 50(1), 151–158. doi:10.1038/s41588-017-0004-9

Li, Y. I., van de Geijn, B., Raj, A., Knowles, D. A., Petti, A. A., Golan, D., … Pritchard, J. K. (2016). RNA splicing is a primary link between genetic variation and disease. Science, 352(6285), 600–604. doi:10.1126/science.aad9417

Liu, L., Sun, L., Li, Z. H., Li, H. M., Wei, L. P., Wang, Y. F., & Qian, Q. J. (2013). BAIAP2 exhibits association to childhood ADHD especially predominantly inattentive subtype in Chinese Han subjects. Behav Brain Funct, 9, 48. doi:10.1186/1744-9081-9-48

Luo, R., Sanders, S. J., Tian, Y., Voineagu, I., Huang, N., Chu, S. H., … Geschwind, D. H. (2012). Genome-wide transcriptome profiling reveals the functional impact of rare de novo and recurrent CNVs in autism spectrum disorders. Am J Hum Genet, 91(1), 38–55. doi:10.1016/j.ajhg.2012.05.011

Macosko, E. Z., Basu, A., Satija, R., Nemesh, J., Shekhar, K., Goldman, M., … McCarroll, S. A. (2015). Highly Parallel Genome-wide Expression Profiling of Individual Cells Using Nanoliter Droplets. Cell, 161(5), 1202–1214. doi:10.1016/j.cell.2015.05.002

Mancarci, B. O., Toker, L., Tripathy, S. J., Li, B., Rocco, B., Sibille, E., & Pavlidis, P. (2017). Cross-Laboratory Analysis of Brain Cell Type Transcriptomes with Applications to Interpretation of Bulk Tissue Data. eNeuro, 4(6). doi:10.1523/ENEURO.0212-17.2017

Mancuso, N., Kichaev, G., Shi, H., Freund, M., Giambartolomei, C., Gusev, A., & Pasaniuc, B. (2017). Probabilistic fine-mapping of transcriptome-wide association studies. bioRxiv. doi:10.1101/236869

Maurano, M. T., Humbert, R., Rynes, E., Thurman, R. E., Haugen, E., Wang, H., … Stamatoyannopoulos, J. A. (2012). Systematic localization of common disease-associated variation in regulatory DNA. Science, 337(6099), 1190–1195. doi:10.1126/science.1222794

McLaren, W., Gil, L., Hunt, S. E., Riat, H. S., Ritchie, G. R., Thormann, A., … Cunningham, F. (2016). The Ensembl Variant Effect Predictor. Genome Biol, 17(1), 122. doi:10.1186/s13059-016-0974-4

Miller, J. A., Ding, S. L., Sunkin, S. M., Smith, K. A., Ng, L., Szafer, A., … Lein, E. S. (2014). Transcriptional landscape of the prenatal human brain. Nature, 508(7495), 199–206. doi:10.1038/nature13185

Mirnics, K., Middleton, F. A., Marquez, A., Lewis, D. A., & Levitt, P. (2000). Molecular characterization of schizophrenia viewed by microarray analysis of gene expression in prefrontal cortex. Neuron, 28(1), 53–67.

Mostafavi, S., Battle, A., Zhu, X., Urban, A. E., Levinson, D., Montgomery, S. B., & Koller, D. (2013). Normalizing RNA-sequencing data by modeling hidden covariates with prior knowledge. PLoS One, 8(7), e68141. doi:10.1371/journal.pone.0068141

Nepusz, G., & Csárdi, G. (2006). The igraph software package for complex network research. Complex Systems, 1695(5), 1–9.

Nica, A. C., & Dermitzakis, E. T. (2013). Expression quantitative trait loci: present and future. Philos Trans R Soc Lond B Biol Sci, 368(1620), 20120362. doi:10.1098/rstb.2012.0362

Nica, A. C., Parts, L., Glass, D., Nisbet, J., Barrett, A., Sekowska, M., … Mu, T. C. (2011). The architecture of gene regulatory variation across multiple human tissues: the MuTHER study. PLoS Genet, 7(2), e1002003. doi:10.1371/journal.pgen.1002003

Nord, A. S., Pattabiraman, K., Visel, A., & Rubenstein, J. L. R. (2015). Genomic perspectives of transcriptional regulation in forebrain development. Neuron, 85(1), 27–47. doi:10.1016/j.neuron.2014.11.011

Ojeda, S. R., Hill, J., Hill, D. F., Costa, M. E., Tapia, V., Cornea, A., & Ma, Y. J. (1999). The Oct-2 POU domain gene in the neuroendocrine brain: a transcriptional regulator of mammalian puberty. Endocrinology, 140(8), 3774–3789. doi:10.1210/endo.140.8.6941

Ongen, H., Buil, A., Brown, A. A., Dermitzakis, E. T., & Delaneau, O. (2016). Fast and efficient QTL mapper for thousands of molecular phenotypes. Bioinformatics, 32(10), 1479–1485. doi:10.1093/bioinformatics/btv722

Pagès, H., Carlson, M., Falcon, S., & Li, N. (2018). AnnotationDbi: Annotation Database Interface. R package version 1.42.1.

Pardinas, A. F., Holmans, P., Pocklington, A. J., Escott-Price, V., Ripke, S., Carrera, N., … Consortium, C. (2018). Common schizophrenia alleles are enriched in mutation-intolerant genes and in regions under strong background selection. Nat Genet, 50(3), 381–389. doi:10.1038/s41588-018-0059-2

Parikshak, N. N., Gandal, M. J., & Geschwind, D. H. (2015). Systems biology and gene networks in neurodevelopmental and neurodegenerative disorders. Nat Rev Genet, 16(8), 441–458. doi:10.1038/nrg3934

Parikshak, N. N., Luo, R., Zhang, A., Won, H., Lowe, J. K., Chandran, V., … Geschwind, D. H. (2013). Integrative functional genomic analyses implicate specific molecular pathways and circuits in autism. Cell, 155(5), 1008–1021. doi:10.1016/j.cell.2013.10.031

Polderman, T. J., Benyamin, B., de Leeuw, C. A., Sullivan, P. F., van Bochoven, A., Visscher, P. M., & Posthuma, D. (2015). Meta-analysis of the heritability of human traits based on fifty years of twin studies. Nat Genet, 47(7), 702–709. doi:10.1038/ng.3285

Polioudakis, D., de la Torre-Ubieta, L., Langerman, J., Elkins, A. G., Stein, J. L., Vuong, C. K., … Geschwind, D. H. (2018). A single cell transcriptomic analysis of human neocortical development. bioRxiv. doi:10.1101/401885

Pollen, A. A., Nowakowski, T. J., Chen, J., Retallack, H., Sandoval-Espinosa, C., Nicholas, C. R., … Kriegstein, A. R. (2015). Molecular identity of human outer radial glia during cortical development. Cell, 163(1), 55–67. doi:10.1016/j.cell.2015.09.004

Preciados, M., Yoo, C., & Roy, D. (2016). Estrogenic Endocrine Disrupting Chemicals Influencing NRF1 Regulated Gene Networks in the Development of Complex Human Brain Diseases. Int J Mol Sci, 17(12). doi:10.3390/ijms17122086

Quesnel-Vallieres, M., Irimia, M., Cordes, S. P., & Blencowe, B. J. (2015). Essential roles for the splicing regulator nSR100/SRRM4 during nervous system development. Genes Dev, 29(7), 746–759. doi:10.1101/gad.256115.114

Rakic, P. (1995). A small step for the cell, a giant leap for mankind: a hypothesis of neocortical expansion during evolution. Trends Neurosci, 18(9), 383–388.

Rakic, P. (2009). Evolution of the neocortex: a perspective from developmental biology. Nat Rev Neurosci, 10(10), 724–735. doi:10.1038/nrn2719

Ramasamy, A., Trabzuni, D., Guelfi, S., Varghese, V., Smith, C., Walker, R., … Weale, M. E. (2014). Genetic variability in the regulation of gene expression in ten regions of the human brain. Nat Neurosci, 17(10), 1418–1428. doi:10.1038/nn.3801

Ripke, S., O’Dushlaine, C., Chambert, K., Moran, J. L., Kahler, A. K., Akterin, S., … Sullivan, P. F. (2013). Genome-wide association analysis identifies 13 new risk loci for schizophrenia. Nat Genet, 45(10), 1150–1159. doi:10.1038/ng.2742

Roadmap Epigenomics, C., Kundaje, A., Meuleman, W., Ernst, J., Bilenky, M., Yen, A., … Kellis, M. (2015). Integrative analysis of 111 reference human epigenomes. Nature, 518(7539), 317–330. doi:10.1038/nature14248

Ruzzo, E. K., Perez-Cano, L., Jung, J.-Y., Wang, L.-k., Kashef-Haghighi, D., Hartl, C., … Wall, D. P. (2018). Whole genome sequencing in multiplex families reveals novel inherited and de novo genetic risk in autism. bioRxiv. doi:10.1101/338855

Samocha, K. E., Robinson, E. B., Sanders, S. J., Stevens, C., Sabo, A., McGrath, L. M., … Daly, M. J. (2014). A framework for the interpretation of de novo mutation in human disease. Nat Genet, 46(9), 944–950. doi:10.1038/ng.3050

Sanders, S. J., Murtha, M. T., Gupta, A. R., Murdoch, J. D., Raubeson, M. J., Willsey, A. J., … State, M. W. (2012). De novo mutations revealed by whole-exome sequencing are strongly associated with autism. Nature, 485(7397), 237–241. doi:10.1038/nature10945

Schaid, D. J., Chen, W., & Larson, N. B. (2018). From genome-wide associations to candidate causal variants by statistical fine-mapping. Nat Rev Genet, 19(8), 491–504. doi:10.1038/s41576-018-0016-z

Schaub, M. A., Boyle, A. P., Kundaje, A., Batzoglou, S., & Snyder, M. (2012). Linking disease associations with regulatory information in the human genome. Genome Res, 22(9), 1748–1759. doi:10.1101/gr.136127.111

Schizophrenia Working Group of the Psychiatric Genomics, C. (2014). Biological insights from 108 schizophrenia-associated genetic loci. Nature, 511(7510), 421–427. doi:10.1038/nature13595

Schmidt, E. M., Zhang, J., Zhou, W., Chen, J., Mohlke, K. L., Chen, Y. E., & Willer, C. J. (2015). GREGOR: evaluating global enrichment of trait-associated variants in epigenomic features using a systematic, data-driven approach. Bioinformatics, 31(16), 2601–2606. doi:10.1093/bioinformatics/btv201

Shabalin, A. A. (2012). Matrix eQTL: ultra fast eQTL analysis via large matrix operations. Bioinformatics, 28(10), 1353–1358. doi:10.1093/bioinformatics/bts163

Shinawi, M., Liu, P., Kang, S. H., Shen, J., Belmont, J. W., Scott, D. A., … Lupski, J. R. (2010). Recurrent reciprocal 16p11.2 rearrangements associated with global developmental delay, behavioural problems, dysmorphism, epilepsy, and abnormal head size. J Med Genet, 47(5), 332–341. doi:10.1136/jmg.2009.073015

Siepel, A., Bejerano, G., Pedersen, J. S., Hinrichs, A. S., Hou, M., Rosenbloom, K., … Haussler, D. (2005). Evolutionarily conserved elements in vertebrate, insect, worm, and yeast genomes. Genome Res, 15(8), 1034–1050. doi:10.1101/gr.3715005

Silbereis, J. C., Pochareddy, S., Zhu, Y., Li, M., & Sestan, N. (2016). The Cellular and Molecular Landscapes of the Developing Human Central Nervous System. Neuron, 89(2), 248–268. doi:10.1016/j.neuron.2015.12.008

Storey, J. D., & Tibshirani, R. (2003). Statistical significance for genomewide studies. Proc Natl Acad Sci U S A, 100(16), 9440–9445. doi:10.1073/pnas.1530509100

Stranger, B. E., Nica, A. C., Forrest, M. S., Dimas, A., Bird, C. P., Beazley, C., … Dermitzakis, E. T. (2007). Population genomics of human gene expression. Nat Genet, 39(10), 1217–1224. doi:10.1038/ng2142

Strunz, T., Grassmann, F., Gayan, J., Nahkuri, S., Souza-Costa, D., Maugeais, C., … Weber, B. H. F. (2018). A mega-analysis of expression quantitative trait loci (eQTL) provides insight into the regulatory architecture of gene expression variation in liver. Sci Rep, 8(1), 5865. doi:10.1038/s41598-018-24219-z

Suls, A., Jaehn, J. A., Kecskes, A., Weber, Y., Weckhuysen, S., Craiu, D. C., … Euro, E. R. E. S. C. (2013). De novo loss-of-function mutations in CHD2 cause a fever-sensitive myoclonic epileptic encephalopathy sharing features with Dravet syndrome. Am J Hum Genet, 93(5), 967–975. doi:10.1016/j.ajhg.2013.09.017

Sunkin, S. M., Ng, L., Lau, C., Dolbeare, T., Gilbert, T. L., Thompson, C. L., … Dang, C. (2013). Allen Brain Atlas: an integrated spatio-temporal portal for exploring the central nervous system. Nucleic Acids Res, 41(Database issue), D996–D1008. doi:10.1093/nar/gks1042

Takata, A., Matsumoto, N., & Kato, T. (2017). Genome-wide identification of splicing QTLs in the human brain and their enrichment among schizophrenia-associated loci. Nat Commun, 8, 14519. doi:10.1038/ncomms14519

Tasic, B., Menon, V., Nguyen, T. N., Kim, T. K., Jarsky, T., Yao, Z., … Zeng, H. (2016). Adult mouse cortical cell taxonomy revealed by single cell transcriptomics. Nat Neurosci, 19(2), 335–346. doi:10.1038/nn.4216

Teyssier, J. R., Rey, R., Ragot, S., Chauvet-Gelinier, J. C., & Bonin, B. (2013). Correlative gene expression pattern linking RNF123 to cellular stress-senescence genes in patients with depressive disorder: implication of DRD1 in the cerebral cortex. J Affect Disord, 151(2), 432–438. doi:10.1016/j.jad.2013.04.010

Toma, C., Hervas, A., Balmana, N., Vilella, E., Aguilera, F., Cusco, I., … Bayes, M. (2011). Association study of six candidate genes asymmetrically expressed in the two cerebral hemispheres suggests the involvement of BAIAP2 in autism. J Psychiatr Res, 45(2), 280–282. doi:10.1016/j.jpsychires.2010.09.001

Turner, S. D. (2014). qqman: an R package for visualizing GWAS results using Q-Q and manhattan plots. bioRxiv. doi:10.1101/005165

Turner, T. N., Yi, Q., Krumm, N., Huddleston, J., Hoekzema, K., Ha, F. S., … Eichler, E. E. (2017). denovo-db: a compendium of human de novo variants. Nucleic Acids Res, 45(D1), D804–D811. doi:10.1093/nar/gkw865

Van der Auwera, G. A., Carneiro, M. O., Hartl, C., Poplin, R., Del Angel, G., Levy-Moonshine, A., … DePristo, M. A. (2013). From FastQ data to high confidence variant calls: the Genome Analysis Toolkit best practices pipeline. Curr Protoc Bioinformatics, 43, 11 10 11–33. doi:10.1002/0471250953.bi1110s43

Veerappa, A. M., Saldanha, M., Padakannaya, P., & Ramachandra, N. B. (2014). Family based genome-wide copy number scan identifies complex rearrangements at 17q21.31 in dyslexics. Am J Med Genet B Neuropsychiatr Genet, 165B(7), 572–580. doi:10.1002/ajmg.b.32260

Veyrieras, J. B., Kudaravalli, S., Kim, S. Y., Dermitzakis, E. T., Gilad, Y., Stephens, M., & Pritchard, J. K. (2008). High-resolution mapping of expression-QTLs yields insight into human gene regulation. PLoS Genet, 4(10), e1000214. doi:10.1371/journal.pgen.1000214

Visel, A., Rubin, E. M., & Pennacchio, L. A. (2009). Genomic views of distant-acting enhancers. Nature, 461(7261), 199–205. doi:10.1038/nature08451

Wainberg, M., Sinnott-Armstrong, N., Knowles, D., Golan, D., Ermel, R., Ruusalepp, A., … Kundaje, A. (2017). Vulnerabilities of transcriptome-wide association studies. bioRxiv. doi:10.1101/206961

Wang, E., Dimova, N., & Cambi, F. (2007). PLP/DM20 ratio is regulated by hnRNPH and F and a novel G-rich enhancer in oligodendrocytes. Nucleic Acids Res, 35(12), 4164–4178. doi:10.1093/nar/gkm387

Ward, L. D., & Kellis, M. (2012). Interpreting noncoding genetic variation in complex traits and human disease. Nat Biotechnol, 30(11), 1095–1106. doi:10.1038/nbt.2422

Wei, T., & Simko, V. (2017). corrplot: Visualization of a Correlation Matrix (Version 0.84). R package version 1.42.1.

Weinberger, D. R. (1987). Implications of normal brain development for the pathogenesis of schizophrenia. Arch Gen Psychiatry, 44(7), 660–669.

Weiss, L. A., Shen, Y., Korn, J. M., Arking, D. E., Miller, D. T., Fossdal, R., … Autism, C. (2008). Association between microdeletion and microduplication at 16p11.2 and autism. N Engl J Med, 358(7), 667–675. doi:10.1056/NEJMoa075974

Willer, C. J., Li, Y., & Abecasis, G. R. (2010). METAL: fast and efficient meta-analysis of genomewide association scans. Bioinformatics, 26(17), 2190–2191. doi:10.1093/bioinformatics/btq340

Willsey, A. J., Sanders, S. J., Li, M., Dong, S., Tebbenkamp, A. T., Muhle, R. A., … State, M. W. (2013). Coexpression networks implicate human midfetal deep cortical projection neurons in the pathogenesis of autism. Cell, 155(5), 997–1007. doi:10.1016/j.cell.2013.10.020

Winden, K. D., Oldham, M. C., Mirnics, K., Ebert, P. J., Swan, C. H., Levitt, P., … Geschwind, D. H. (2009). The organization of the transcriptional network in specific neuronal classes. Mol Syst Biol, 5, 291. doi:10.1038/msb.2009.46

Won, H., de la Torre-Ubieta, L., Stein, J. L., Parikshak, N. N., Huang, J., Opland, C. K., … Geschwind, D. H. (2016). Chromosome conformation elucidates regulatory relationships in developing human brain. Nature, 538(7626), 523–527. doi:10.1038/nature19847

Wu, Q., & Maniatis, T. (1999). A striking organization of a large family of human neural cadherin-like cell adhesion genes. Cell, 97(6), 779–790.

Yang, J., Lee, S. H., Goddard, M. E., & Visscher, P. M. (2011). GCTA: a tool for genome-wide complex trait analysis. Am J Hum Genet, 88(1), 76–82. doi:10.1016/j.ajhg.2010.11.011

Yang, Y. C., Di, C., Hu, B., Zhou, M., Liu, Y., Song, N., … Lu, Z. J. (2015). CLIPdb: a CLIP-seq database for protein-RNA interactions. BMC Genomics, 16, 51. doi:10.1186/s12864-015-1273-2

Zhang, B., & Horvath, S. (2005). A general framework for weighted gene co-expression network analysis. Stat Appl Genet Mol Biol, 4, Article17. doi:10.2202/1544-6115.1128

Zhang, F., Wang, G., Shugart, Y. Y., Xu, Y., Liu, C., Wang, L., … Zhang, D. (2014). Association analysis of a functional variant in ATXN2 with schizophrenia. Neurosci Lett, 562, 24–27. doi:10.1016/j.neulet.2013.12.001

Zhang, Y., Chen, K., Sloan, S. A., Bennett, M. L., Scholze, A. R., O’Keeffe, S., … Wu, J. Q. (2014). An RNA-sequencing transcriptome and splicing database of glia, neurons, and vascular cells of the cerebral cortex. J Neurosci, 34(36), 11929–11947. doi:10.1523/JNEUROSCI.1860-14.2014

Zhang, Y., Sloan, S. A., Clarke, L. E., Caneda, C., Plaza, C. A., Blumenthal, P. D., … Barres, B. A. (2016). Purification and Characterization of Progenitor and Mature Human Astrocytes Reveals Transcriptional and Functional Differences with Mouse. Neuron, 89(1), 37–53. doi:10.1016/j.neuron.2015.11.013

Zhou, W., Shi, Y., Li, F., Wu, X., Huai, C., Shen, L., … Qin, S. (2018). Study of the association between Schizophrenia and microduplication at the 16p11.2 locus in the Han Chinese population. Psychiatry Res, 265, 198–199. doi:10.1016/j.psychres.2018.04.049

Zhou, X., Carbonetto, P., & Stephens, M. (2013). Polygenic modeling with bayesian sparse linear mixed models. PLoS Genet, 9(2), e1003264. doi:10.1371/journal.pgen.1003264

